# The npBAF to nBAF Chromatin Switch Regulates Cell Cycle Exit in the Developing Mammalian Cortex

**DOI:** 10.1101/2020.01.17.910794

**Authors:** Simon MG Braun, Ralitsa Petrova, Jiong Tang, Andrey Krokhotin, Erik L Miller, Yitai Tang, Georgia Panagiotakos, Gerald R Crabtree

## Abstract

Nervous system development is orchestrated by tightly-regulated progenitor cell divisions, followed by differentiation at precise but varying times across different regions. As progenitors exit the cell cycle, they initiate a subunit switch of the mSWI/SNF or npBAF complex to generate neuron-specific nBAF complexes. These chromatin regulatory complexes play dosage-sensitive roles in neural development and are frequently mutated in neurodevelopmental disorders. Here we manipulated the timing of BAF subunit exchange in the developing mouse brain and find that deletion of the npBAF subunit BAF53a blocks progenitor proliferation, leading to impaired neurogenesis. We show that npBAF complexes regulate cell cycle progression by antagonizing Polycomb complexes to promote chromatin accessibility at cell cycle and NPC identity genes. Replacement of the actin-related protein, Actl6a (BAF53a) by the neuron-specific actin-related protein, Actl6b (BAF53b), but not other regulators of proliferation, rescues progenitors by promoting neuronal differentiation. We propose that the function of the npBAF to nBAF chromatin switch is to control progenitor cell cycle exit and promote synchronous neural differentiation.

## INTRODUCTION

Neurogenesis in vertebrates is accompanied by dramatic alterations in chromatin structure that correlate with temporally precise developmental transitions. The discovery that many human neurologic diseases have their origin in mutations in chromatin regulators has called attention to the different roles played by these chromatin regulatory mechanisms. One event that correlates with cell cycle exit in the mammalian nervous system is a switch in subunit composition of BAF or mSWI/SNF complexes. These ATP-dependent chromatin regulators are similar to yeast SWI/SNF and Drosophila BAP complexes, and in mammals are comprised of 15 subunits encoded by 29 genes that are combinatorially assembled to produce several hundred complexes ^1-7^. BAF subunits are exchanged at the transitions from pluripotent stem cells to neural progenitor cells to neurons, resulting in highly specific neuronal nBAF complexes ^8^. As neural progenitor cells (NPCs) in the developing nervous system exit the cell cycle, neural progenitor BAF (npBAF) complexes containing BAF53a (*Actl6a*), BAF45a/d and SS18 switch subunits to give rise to nBAF complexes containing the neuron-specific subunits BAF53b (*Actl6b*), BAF45b/c and CREST ^5^. Biochemical analysis and single cell RNA sequencing studies indicate that a single neuron likely expresses several hundred complexes of distinct subunit composition ^9^. These complexes are highly stable *in vitro*, allowing for the generation of distinct composite surfaces, much like letters in words ^10^, thought to be responsible for their wide range of biologically specific activities ^11^.

Consistent with the diverse roles played by these complexes during neural development, mutations in BAF subunits are frequently associated with neurodevelopmental diseases, including autism spectrum disorders (ASD), intellectual disability (ID), and speech disorders. Mutations in *SMARCB1* (encoding BAF47), *ARID1a/b* (encoding Baf250a/b) or *SMARCA4* (encoding Brg1/Brm) cause Coffin-Siris and Nicolaides-Baraitser syndromes ^12-16^, while mutations in *ACTL6B* (encoding BAF53b) cause ASD ^17^. Finally, GWAS studies have implicated sequence variation in *PBRM1* (encoding BAF180) in exceptional intellectual ability ^18,19^. The BAF subunits mutated in human neurodevelopmental and psychiatric disorders identified to date include: BAF250b/*ARID1B*, BAF250a/*ARID1A*, BAF200/*ARID2*, BAF180/*PBRM1*, BCL11A/*BCL11A*, BRG1/*SMARCA4*, BRG2/*SMARCA2*, BAF170/*SMARCC2*, BAF47/*SMARCB1*, BAF57/*SMARCE1*, and BAF45d/*DPF2* ^20,21^. In general, the mutations are heterozygous loss-of-function alleles, indicating a dosage-sensitive role for BAF subunits in the development of the human nervous system.

Cell cycle exit in the developing brain is a highly orchestrated process that appears to function more like an “AND” gate, integrating numerous pathways rather than operating under the control of a single master regulator. A variety of intrinsic cues and extrinsic signals, including depolarization, voltage-gated channels, neurotransmitter receptors, mitotic spindle orientation, growth factor signaling pathways, cellular metabolism and intrinsic transcriptional mechanisms, contribute to cell cycle exit in NPCs ^22-25^. It remains unclear, however, how these mechanisms are coordinated and synchronized with the onset of neural differentiation. In the *Drosophila* nervous system, the Osa subunit of the homologous BAP complex has been shown to regulate the progression of neural progenitors through distinct transcriptional states to control the number of transit-amplifying divisions ^26^.

In mammals, exchange of BAF subunits at the transition from cycling progenitor to post-mitotic neuron is driven by a triple negative genetic circuit — BAF53a expression is suppressed by miR-9 and miR-124, which target the 3’ untranslated region (UTR) of *Baf53a* and are in turn repressed by REST ^6^. Thus, when REST expression is suppressed, miR-9 and miR-124 are activated to lead to repression of BAF53a. Extending expression of the npBAF subunit BAF53a by deleting the miR-9 and miR-124 binding sites from its 3’ UTR was shown to result in excessive progenitor proliferation in the neural tube ^6^, a phenotype resembling the extended expression of another npBAF subunit, BAF45a, in the developing spinal cord ^5^. Moreover, executing this microRNA/chromatin switch to drive the expression of nBAF subunits in human fibroblasts effectively converts these cells into neurons ^9^, highlighting the importance of the BAF complexes in both cell cycle progression and neuronal differentiation. This finding is consistent with observations that BAF53b and CREST mutant mice have normal progenitor proliferation and instead display phenotypes associated with neuronal differentiation and maturation, including impaired activity-dependent dendritic morphogenesis and synaptic plasticity ^4,27,28^. Nevertheless, while the subunit exchange between npBAF and nBAF temporally corresponds to cell cycle exit, the precise role of suppression of the npBAF subunits and replacement by nBAF subunits, as well as the mechanisms by which this switch is involved in the regulation of cell cycle exit, have not been explored.

Here we investigated the role of the npBAF to nBAF switch in cell cycle exit in the embryonic mouse brain. Conditional deletion of the npBAF subunit Actl6a, an actin-related protein, leads to loss of both Actl6a and β-actin from BAF complexes, resulting in a cell cycle block in NPCs and impaired cortical neurogenesis. We show that BAF53a-containing npBAF complexes are required to promote chromatin accessibility at a broad array of cell cycle and NPC identity genes through opposition of Polycomb repressive complexes. Importantly, we find that the cell cycle block that results from *Baf53a* deletion can be rescued by expression of neuron-specific nBAF subunits, allowing for cell cycle exit and neuronal differentiation. We propose that the switch from npBAF to nBAF chromatin regulators functions as a checkpoint allowing for the faithful execution of differentiation programs in the developing cortex.

## RESULTS

### BAF53a regulates cell cycle progression in the developing cortex

During brain development, the exit of NPCs from the cell cycle and subsequent onset of neuronal differentiation are closely correlated with repression of BAF53a and activation of BAF53b. Consistent with this, we observed that BAF53a is enriched in SOX2-expressing radial glia and TBR2-expressing intermediate progenitor cells (IPCs) in the ventricular and subventricular zones of the mouse embryonic cortex (VZ and SVZ, respectively). In contrast, BAF53b is highly expressed in MAP2ab+ neurons in the cortical plate (CP) (**Fig. 1a**). Similarly, analysis of published single cell RNA-seq data ^29^ from the developing mouse cortex at embryonic day 12 (E12) to E15 confirmed that npBAF subunit expression (BAF53a, BAF45d and SS18) is enriched in NPCs in the VZ and SVZ, and nBAF subunits (BAF53b, BAF45b and CREST) are expressed in newly generated cortical neurons (**Supplementary Fig. 1a**).

**Figure 1.**
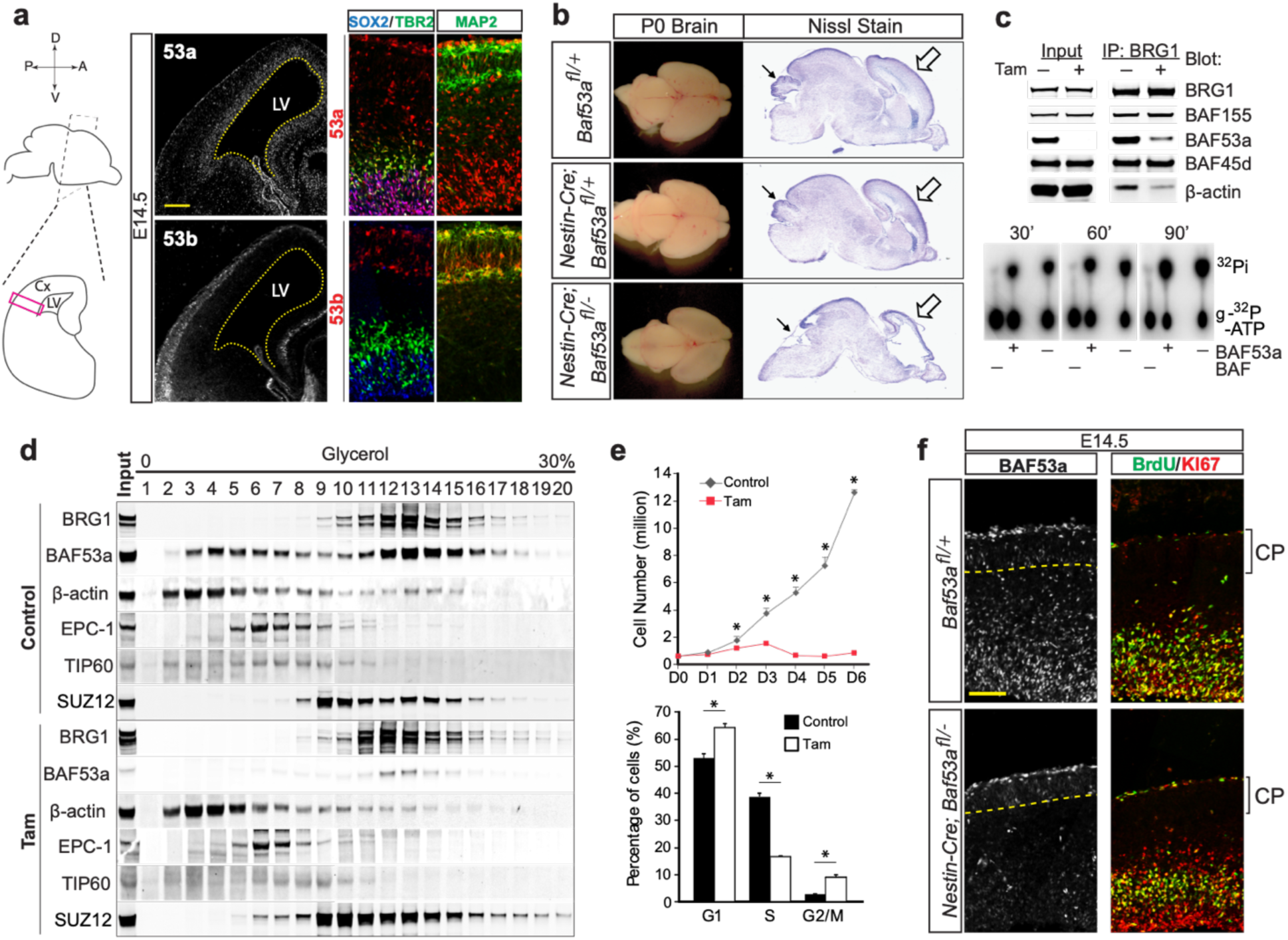
BAF53a regulates cell cycle progression in the developing cortex. **(a)** Representative coronal sections through the developing mouse brain at E14.5 depicting expression of BAF53a (*top left*) and BAF53b (*bottom left*). Dashed line delineates the wall of the lateral ventricle (LV). Immunofluorescence staining shows BAF53A expression in SOX2+ (*blue*) and TBR2+ (*green*) neural stem and intermediate progenitor cells, respectively (*top right*). Immunolabeling for BAF53b and MAP2 (green, *bottom right*) reveals strong BAF53b expression in post-mitotic neurons in the cortical plate at E14.5. *Scale bars, 50μm*. **(b)** BAF53a mutant brains at P0. Photomicrographs of whole control, heterozygous and BAF53a mutant brains at P0 (*left*) and Nissl stains of sagittal sections (*right*). Closed arrow points to cerebellum and open arrow indicates the neocortex. **(c)** Immunoprecipitation and western blot analysis of BAF proteins in control and *Baf53a* mutant nuclear extracts using an anti-BRG1 antibody (*top panel*). γ-32P labeled-ATP was used as substrate to assess BAF ATPase activity, separated by chromatography and visualized by autoradiography (*bottom panel*). **(d)** Western blot on BAF53a-deficient and control nuclear glycerol gradients (0-30 %) fractions analyzed for BAF, PRC2 and TIP60 complex components. **(e)** Growth curve of *Actin-CreER*; *Baf53a*^*fl/–*^ neurospheres for 6 days after Tamoxifen (Tam) or Ethanol (EtOH) control treatment (*top panel*). Quantification of cell cycle FACS analysis of *Actin-CreER*; *Baf53a*^*fl/–*^ neurospheres. Cells were treated with BrdU 2 hours before collection and harvested 72 hours after tamoxifen administration for FACS (*bottom panel*). n=3, error bars represent SEM, *p<0.05. **(f)** Immunohistochemistry of E14.5 mouse cortex showing reduced BAF53a expression in the ventricular zone of *Nestin-Cre;Baf53a* ^*fl/–*^ brains compared to controls (*left panel*). Immunostaining for the proliferative marker KI67 (*red*) and short-term BrdU labeling (*green*, 3.5h pulse) (*right panel*). *Scale bar, 50μm*.

To investigate whether the npBAF subunit BAF53a is required for cell cycle progression in the nervous system, we employed a conditional genetic approach to delete *Baf53a*. As constitutive deletion of *Baf53a* leads to early embryonic lethality (data not shown), we generated *Baf53a* conditional mutant mice in which exons 4-5 are flanked by loxP sites, *Baf53a*^fl/fl^ (**Supplementary Fig. 1b**). *Baf53a*^fl/-^ mice were then crossed with *Nestin-Cre* mice to inactivate BAF53a in NPCs throughout the brain at embryonic day 10.5, before its normal developmental repression. *Nestin-Cre;Baf53a*^fl*/-*^ (BAF53a mutant) mice survived to birth, but died perinatally due to a failure to nurse. The expression pattern of BAF53b was not significantly altered in BAF53a mutant mice (**Supplementary Fig. 1c**). BAF53a mutants were severely microcephalic with a dramatic reduction of cortical thickness, enlarged lateral ventricles, and a hypoplastic cerebellum (**Fig. 1b, Supplementary Fig. 1d**), suggesting a direct role for BAF53a in the control of brain size.

To assess BAF complex assembly and function biochemically, we bred conditional *Baf53a* mutant mice to a transgenic mouse line expressing a tamoxifen-inducible form of Cre recombinase (Cre-ER) ^30^ under the control of the Actin promoter (*Actin-CreER;Baf53a*^fl/-^). We then cultured E12.5 NPCs isolated from *Actin-CreER;Baf53a*^fl/-^ mice *in vitro*. Following tamoxifen treatment, mutant BAF complexes continued to assemble into large multi-subunit chromatin remodeling complexes, as shown by immunoprecipitation for the core BAF complex subunit BRG1 and sucrose gradient sedimentation analysis (**Fig. 1c, 1d**). We found that complexes containing BRG1 and other npBAF subunits, but lacking BAF53a, lacked β-actin but maintained normal DNA-dependent ATPase activity (**Fig. 1c, 1d, Supplementary Fig. 2a**), suggesting that BAF53a and β-actin are not required for BAF complex assembly and function. The loss of these subunits also did not alter global chromatin affinity of the BAF complex in NPCs, as measured by extracting chromatin-associated proteins upon treating nuclei with increasing amounts of salt (**Supplementary Fig. 2b**).

The proliferation of NPCs lacking BAF53a, however, was severely impaired in culture, with mutant cells stalled at G1 and G2/M phases of the cell cycle (**Fig. 1e**). In support of a mechanism of cell cycle arrest, we found that deletion of *Baf53a* results in a significant increase in anaphase bridges (**Supplementary Fig. 2c**), a hallmark of improper segregation of daughter chromosomes during mitosis. In previous studies in ES cells, we and others have found that anaphase bridges detected after *Brg1* subunit deletion were the result of a failure of Topoisomerase II to bind to DNA and relieve catenated chromosomes ^31^. Cells with anaphase bridges activate the decatenation checkpoint through the action of the ATR/ATM kinases and stall progress through the cell cycle. Unless this checkpoint blockade is relieved, they undergo apoptosis ^32^. Consistent with the idea of decatenation checkpoint-induced cell death, we also observed an increase in cell death in cultured NPCs following *Baf53a* deletion (**Supplementary Fig. 2d**). Using a second, complementary strategy, we introduced lentiviruses expressing miR-9 and miR-124 into NPCs to induce early reduction in BAF53a expression *in vitro*. This dramatically reduced BAF53a levels and produced a prominent G2/M cell cycle arrest similar to that seen in BAF53a mutant cells (**Supplementary Fig. 2e, 2f**). In line with these *in vitro* findings, following a 3-hour pulse of the thymidine analog bromodeoxyuridine (BrdU) *in vivo*, we observed a reduction in BrdU+/KI67+ cells in the VZ of BAF53a mutant mice at E14.5 and E15.5 (**Fig. 1f, Supplementary Fig. 1e, 1f**). Together these data suggest that early ablation of BAF53a results in a defect in cell cycle progression and NPC proliferation in part due to the decatenation checkpoint.

### B*af53a* deletion impairs the maintenance of neural stem and progenitor cells in the cortical germinal zones

To characterize the consequences of early ablation of BAF53a on cortical neurogenesis, we performed immunofluorescence staining for the proliferation marker KI67 and the mitotic indicator phosphorylated histone H3 (PH3) (**Fig. 2a, 2b, Supplementary Fig. 3a)**. PH3+ cells are normally observed adjacent to the ventricular wall and scattered in the SVZ in control mice (**Fig. 2b**). In BAF53a mutant mice at E15.5, we observed increased numbers of KI67+ cells, in addition to accumulation of large, round PH3+ and KI67+ cells in VZ and SVZ, further underscoring a potential block in the G2/M phase of the cell cycle (**Fig. 2a, 2b, Supplementary Fig. 3a)**. This increase in PH3+ cells persisted at later developmental stages (**Supplementary Fig. 3c**) and was accompanied by an increase in cleaved caspase 3 staining, indicating an increase in cell death upon *Baf53a* deletion (**Fig. 2a, 2b, Supplementary Fig. 3b**) that is consistent with certain cells failing the decatenation checkpoint. Strikingly, immunostaining for neural stem and progenitor cell markers also revealed a major disorganization of the germinal zones in BAF53a mutants. In control mice, PAX6 and TBR2 label discrete populations of VZ apical progenitors and SVZ intermediate progenitors (IPCs), respectively. However, in BAF53a mutant mice, the progenitor pools appear mixed (**Fig. 2c**), with displacement of radial glial cells away from the ventricular wall and accumulation of TBR2+ IPCs closer to the ventricle. In addition, we observed an increased number of cells co-expressing PAX6 and TBR2, indicative of an abnormal mixing of cell identity markers in BAF53a knockout NPC populations. Immunostaining for the radial glial cell marker phospho-Vimentin (pVim) further illuminated a disruption of the neurogenic niche in BAF53a mutant mice (**Fig. 2d**), as we observed increased numbers of pVim+ cells in the SVZ away from the ventricle. In the mutant brains, both pVim+ radial glia and TBR2+ IPCs also expressed high levels of H2AX (**Fig. 2d**), indicating an accumulation of DNA double stranded breaks, consistent with the increased number of anaphase bridges we observed *in vitro* (**Supplementary Fig. 2c**) and lending additional support to checkpoint arrest in BAF53a mutant NPCs.

**Figure 2.**
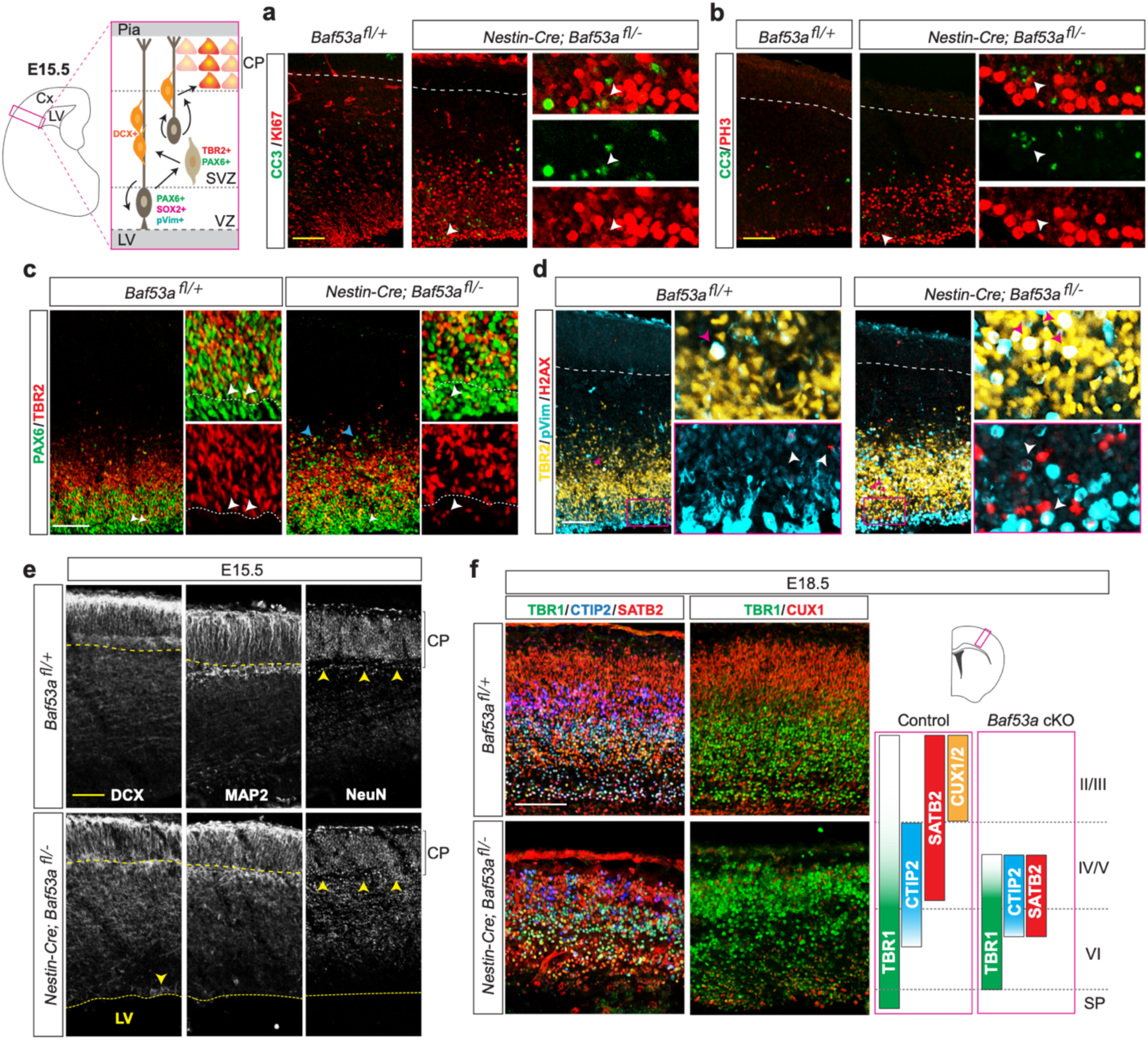
*BAF53a* deletion impairs the maintenance of neural stem and progenitor populations in the germinal zones of the cortex. (**a, b**) Schematic of mouse cortical neurogenesis (*left*) depicting cortical radial glia (*brown*), and their progeny, intermediate progenitor cells (*tan*) and migratory neuroblasts (*orange*), as well as cell type-specific markers expressed by each population. (**a**) Immunostaining for the proliferation marker KI67 (*red*), or (**b**) the M-phase marker phospho-histone H3 (PH3; *red*), showing co-localization with the cell apoptosis marker cleaved caspase 3 (CC3, *green*) in E15.5 BAF53a mutant cortex. *Scale bar, 50μm*. **(c)** Co-labeling for the radial glia and intermediate progenitor markers PAX6 (*green*) and TBR2 (*red*), respectively, depicts disorganization of the stem and progenitor pools in the ventricular and subventricular zones (VZ/SVZ) (*insets*) in E15.5 BAF53a mutant brains. *White arrow* points to TBR2+ cells in the VZ of mutant brains, and *blue* arrow points to the presence of PAX6+ cells further away from the ventricle in mutant mice compared to controls. *Scale bar, 50μm*. **(d)** Immunostaining for the radial glia and intermediate progenitor markers phospho-Vimentin (pVim, *cyan*) and TBR2 (*amber*), respectively, further highlights the disorganization of the VZ/SVZ in E15.5 BAF53a mutant cortex, as well as the presence of many cells in the SVZ co-expressing both markers (*upper panels*). pVim+ cells in the VZ of BAF53a mutants are positive for H2A histone family member X (H2AX, *red*) indicating DNA damage (*bottom panels*). *Scale bar, 50μm*. **(e)** Immunofluorescence staining labeling migratory neuroblasts (doublecortin, DCX) and young and mature neurons (MAP2 and NeuN, respectively) showing mis-localized DCX+ neuroblasts (*bottom left*) and NeuN+ neurons (*bottom right*) in the VZ and IZ, respectively, in BAF53a mutant cortex. *Scale bar, 50μm*. **(f)** E18.5 immunostainings of mouse cortical plate for markers of layer VI (TBR1, *green*), layer IV-V (CTIP2, *blue*; SATB2, *red*), and layer II-III (CUX1, *red*) excitatory neurons reveal disorganization of deep cortical layers and loss of superficial layers in BAF53a mutant compared to control brains. *Scale bar, 50μm*.

The disorganization of the progenitor pools in BAF53a mutant mice was reflected in the CP, where populations of newborn neurons were also disrupted. During normal neurogenesis, the cortical layers are generated following successive rounds of progenitor divisions and the concomitant migration and differentiation of newborn neurons into the CP. Interestingly, we detected an increased number of DCX-expressing immature neurons in the VZ of BAF53a mutant mice, suggesting precocious differentiation in a subset of mutant NPCs (**Fig. 2e**). In control mice at E15.5, the CP was observed as a dense layer of NeuN+ neuronal cell bodies overlying the intermediate zone, whereas in BAF53a mutant mice, the NeuN+ domain appeared expanded, and MAP2ab+ neuronal processes had lost their vertical organization (**Fig. 2e**). These apparent defects in differentiation were further amplified at later timepoints in cortical development. BrdU pulse-chase experiments revealed that the majority of NPCs labelled at E15.5 failed to give rise to upper layer neurons that migrated into the CP by E18.5 (**Supplementary Fig. 3d**). Using immunostaining for various markers of laminar identity at E18.5, we also found that TBR1-, CTIP2- and SATB2-expressing neurons were intermixed and failed to form distinct layers. CUX1-expressing upper layer neurons appeared to be lost altogether (**Fig. 2f**), as were BRN1-expressing neurons (**Supplementary Fig. 3e**), suggesting a near-complete absence of later-generated neuronal cell types. Taken together, this phenotypic analysis reveals a central role for BAF53a in cortical neurogenesis. Early ablation of BAF53a in NPCs leads to deficits in cell cycle progression, premature cell cycle exit, and disruption of the neurogenic niche. These defects in the germinal zones in turn have major consequences on neuronal differentiation, resulting in improper mixing of nascent populations and deficits in the generation of later-born lineages.

### Intrinsic and extrinsic drivers of NPC proliferation cannot rescue cell cycle block resulting from *Baf53a* deletion

To determine if BAF53a operates upstream or downstream of known growth signaling pathways *in vivo*, we first asked whether β-catenin could promote NPC cell cycle progression in the absence of BAF53a. Transgenic expression of constitutively active β-catenin in mice has been shown to promote cell cycle re-entry and proliferation of NPCs. As a result, cortical surface area and overall brain size are increased, indicating that Wnt signaling is a powerful driver of NPC expansion ^22^. However, even when β-catenin is constitutively expressed, NPCs still exit the cell cycle, raising the question of which mechanisms control cell cycle exit under these conditions. Therefore, we bred mice in which the expression of constitutively active β-catenin and deletion of *Baf53a* were both driven by the *Nestin-Cre* transgene, permitting concurrent expression of β-catenin and inactivation of BAF53a in NPCs. As expected, expression of active β-catenin in *Baf53a*^fl/+^ mice resulted in excess NPC proliferation, as measured by KI67 staining, and a markedly enlarged telencephalon at E15.5 (**Fig. 3a, 3b, Supplementary Fig. 4a, 4b**). However, expression of active β-catenin in *Baf53a*^fl/-^ NPCs failed to rescue the cell cycle block resulting from *Baf53a* deletion, as reflected by persistently elevated levels of PH3 staining at E15.5. The double mutant brains remained microcephalic and displayed reduced cortical thickness (**Fig. 3a, 3b, Supplementary Fig. 4a, 4b**).

**Figure 3.**
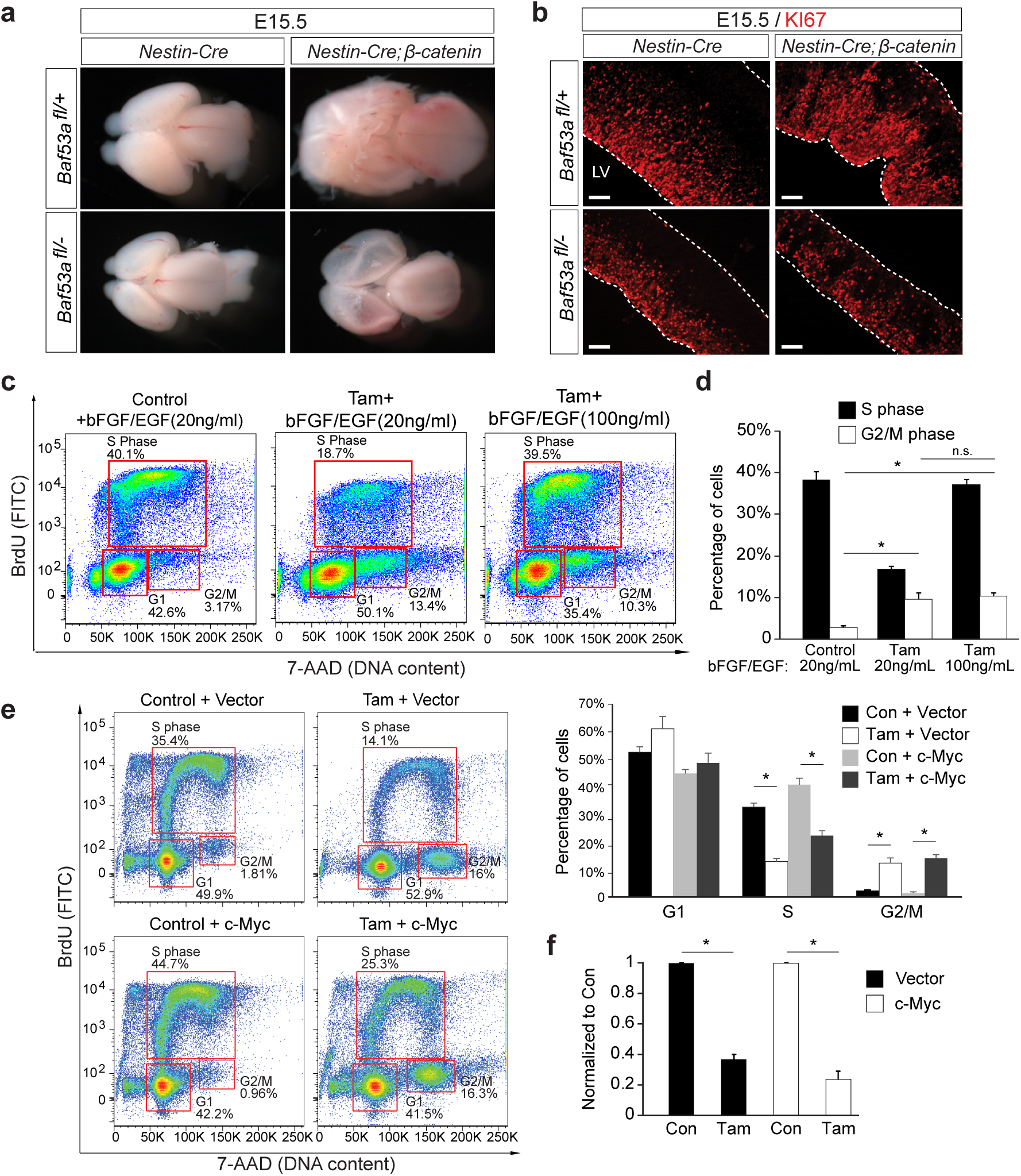
Intrinsic and extrinsic drivers of NPC proliferation cannot rescue proliferation defects resulting from *Baf53a* deletion. **(a)** Photomicrographs of whole brains from BAF53a mutant or control embryos with and without the *β-catenin* transgene at E15.5. **(b)** Immunofluorescence staining for the proliferation marker KI67 on E15.5 sagittal sections through the cortex of BAF53a mutant or control embryos with and without the β-catenin transgene, respectively. *Scale bar, 100μm*. **(c)** Cell cycle FACS analysis of *Actin-CreER*; *Baf53a*^*fl/–*^ neurospheres upon BrdU incorporation with and without tamoxifen (Tam) treatment and following application of increasing concentrations of the growth factors bFGF and EGF. **(d)** Quantification of the mitogens’ effects on cell cycle in the context of *Baf53a* deletion. n=3, error bars represent SEM, *p<0.05. **(e)** Cell cycle FACS analysis depicting the effect of c-*Myc* over-expression in *Actin-CreER*; *Baf53a* ^*fl/–*^ neurospheres with and without tamoxifen. Cells were infected with either empty vector virus or c-*Myc* virus and selected by hygromycin prior to tamoxifen treatment for 72 hours. n=3, error bars represent SEM, *p<0.05. **(f)** Quantification of total cell numbers 72 hours after Tamoxifen treatment (normalized to condition treated with EtOH). n=3, error bars represent SEM, *p<0.05.

Growth factors like FGF2 (bFGF) and EGF have also been shown to promote NPC proliferation ^33^. It is possible that BAF53a-containing BAF complexes control cell cycle progression by driving the production of growth factors or, alternately, that they operate downstream of growth factors, as in the case of β-catenin. We thus examined whether the cell cycle block at G1/S and G2/M observed in BAF53a mutants could be rescued by extrinsic application of growth factors *in vitro*. Specifically, we tested whether increasing concentrations of FGF2 and EGF in the culture medium could compensate for the loss of BAF53a in *Actin-CreER;Baf53a*^fl/-^ NPCs. Under normal proliferative conditions, the presence of EGF and FGF2 in the media (at 20ng/ml) failed to rescue cell cycle block in NPCs lacking BAF53a. Addition of five-fold higher concentrations of these two mitogens could partially rescue the block at G1/S, but did not rescue the G2/M phase block (**Fig. 3c, 3d**). Transduction of NPCs with *c-myc*-overexpressing lentiviruses has also been shown to enhance cell proliferation by promoting G1/S progression ^34-36^. In inducible *Actin-CreER;Baf53a*^fl/-^ NPCs, overexpression of *c-myc* also failed to rescue G2/M block resulting from BAF53a deletion and only partially rescued the block at G1/S (**Fig. 3e, 3f**). In addition, overexpression of *Sox2* and *Pax6*, known regulators of radial glial cell identity and maintenance, increased NPC proliferation in control cultures; however, this was unable to rescue cell cycle block at either G1/S or G2/M in *Baf53a*-deleted NPCs (**Supplementary Fig. 4c, 4d**). Taken together, these studies suggest that BAF53a is necessary for NPC proliferation and that it functions directly on cell cycle regulatory mechanisms downstream of known intrinsic or extrinsic drivers of proliferation.

### BAF53b can rescue the cell cycle block resulting from deletion of BAF53a

As NPCs exit the cell cycle in the developing brain, npBAF complexes containing BAF53a, BAF45a/d and SS18 switch subunits to produce neuron-specific nBAF complexes that contain BAF53b, BAF45b/c and CREST. To determine if BAF subunit exchange has a causal role during neurogenesis, we sought to determine whether BAF53b expression can rescue the cell cycle block and cell death resulting from *Baf53a* deletion in NPCs. We hypothesized that BAF53a mutant NPCs remain blocked in the cell cycle until BAF53b is expressed to form nBAF complexes that promote differentiation. We thus attempted to rescue the cell cycle block in mutant NPCs not by inducing cell cycle progression and proliferation, but rather by promoting cell cycle exit and neural differentiation. To test this hypothesis *in vitro*, we used lentiviruses to overexpress either BAF53a or BAF53b in inducible *Actin-CreER;Baf53a*^fl/-^ NPCs (**Supplementary Fig. 5a**). Following tamoxifen treatment of NPCs and consistent with our previous findings, we observed a cell cycle block at both G1/S and G2/M in cells lacking BAF53a (**Fig. 4a-c**). Restoring BAF53a expression rescued the cell cycle block in mutant cells (**Fig. 4a, 4b**). Strikingly, the cell cycle block in NPCs lacking BAF53a was also rescued by overexpression of BAF53b, suggesting that the switch to nBAF subunits is sufficient to promote cell cycle exit upon repression of BAF53a or that the two proteins have redundant functions (**Fig. 4a-c**). To further explore the fate of to the cells rescued by either BAF53a and BAF53b overexpression, we immunostained for the NPC marker Nestin and the neuronal marker MAP2ab (**Supplementary Fig. 5b, 5c**). As expected, the number of Nestin-expressing cells was dramatically reduced in the BAF53a mutant NPCs. Overexpression of BAF53a or BAF53b was sufficient to rescue Nestin expression (**Supplementary Fig. 5d**), at least in part owing to increased cell survival (**Fig. 4b**). Interestingly, we detected MAP2ab-expressing cells following tamoxifen treatment, even in media containing EGF and bFGF. This suggests that loss of BAF53a is sufficient to promote neuronal differentiation of NPCs and may be required for NPCs to stop responding to extrinsic proliferative signals and initiate differentiation programs. MAP2ab expression further increased with overexpression of BAF53b (**Supplementary Fig. 5d**), in line with the idea that subunit switching enables differentiation onset.

**Figure 4.**
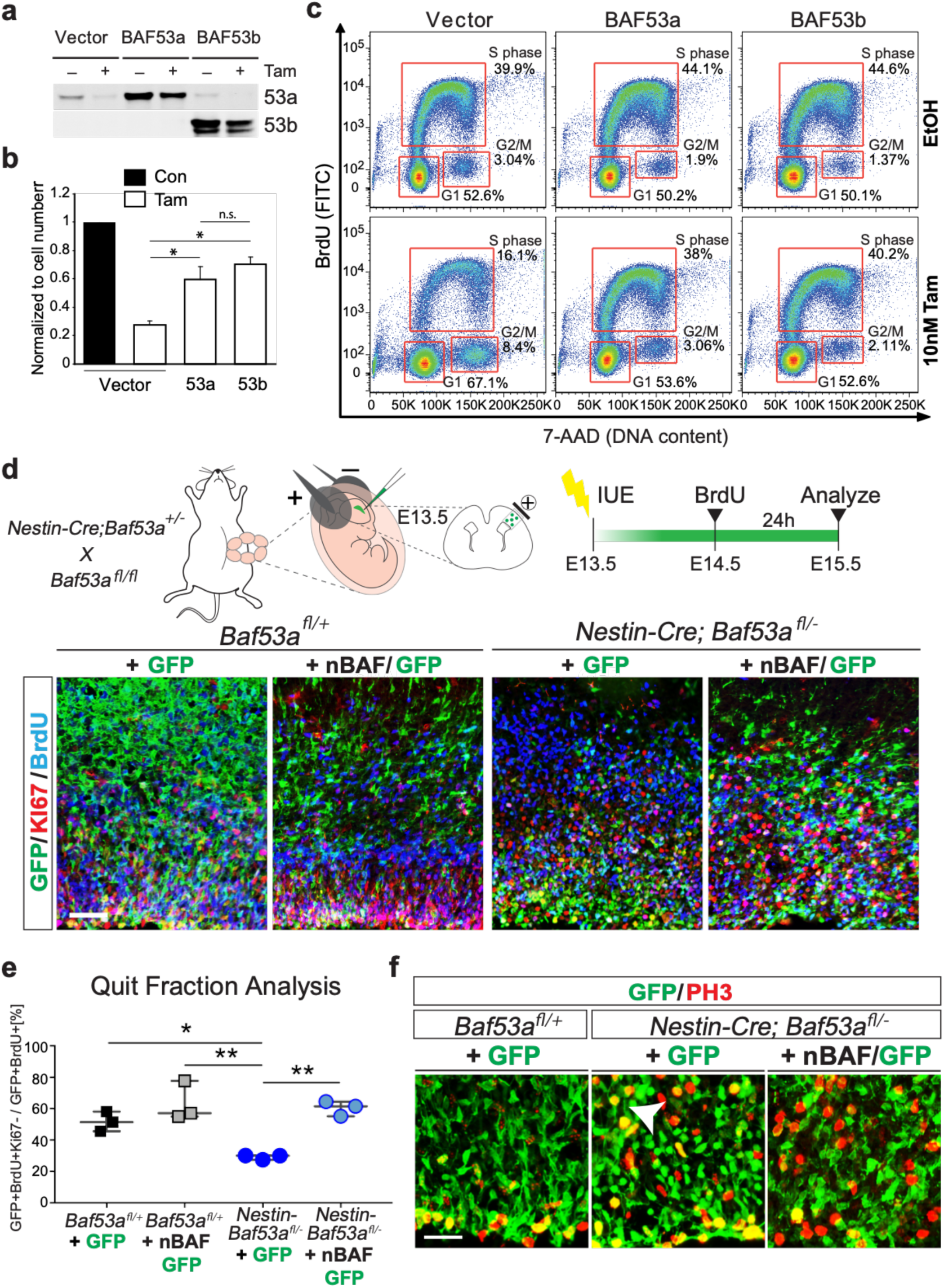
BAF53b can rescue cell cycle block resulting from *Baf53a* deletion *in vitro* and *in vivo*. **(a)** Western blot analysis to measure BAF53a and BAF53b levels in *Actin-CreER*;*Baf53a*^*fl/–*^ neurospheres treated with tamoxifen or vehicle following infection with empty vector or BAF53a or BAF53b overexpressing lentiviruses. **(b)** Quantification of *Actin-CreER*;*Baf53a*^*fl/–*^ total cell numbers 72 hours after Tamoxifen treatment (normalized to control condition infected with empty vector and treated with EtOH). n=3, error bars represent SEM, *p<0.05. **(c)** Cell cycle FACS analysis of *Actin-CreER*;*Baf53a*^*fl/–*^ neurospheres treated with tamoxifen or vehicle (EtOH) infected with empty vector, BAF53a or BAF53b overexpressing lentiviruses. Cells were selected by puromycin prior to tamoxifen treatment. Infected cells were labeled with BrdU for 2 hours prior to FACS analysis 72 hours after tamoxifen administration. **(d)** Schematic of experimental paradigm: in utero electroporation (IUE) of BAF53a mutant and control E13.5 embryos with a control plasmid (GFP), or a mix of GFP and a polycistronic vector co-expressing *Baf53b, CREST, Baf45b* (nBAF). Representative images of coronal sections from E15.5 control *Baf53a*^*fl/+*^ and *Nestin-Cre*; *Baf53a*^*fl/–*^ mutant brains immunostained for GFP (*green*), KI67 (*red*), and BrdU (*blue*). *Scale bars, 50μm*. **(e)** Cell cycle exit analysis of BAF53a mutant and control brains at E15.5 following electroporation with control (GFP) or nBAF vectors (n=3 embryos per condition). Data presented as mean +/- SEM; *\* p<0*.*03, ** p<0*.*004, one-way ANOVA and post-hoc Tukey’s test for multiple comparisons*. **(f)** Immunofluorescence staining of coronal sections through the VZ/SVZ of electroporated BAF53a mutant and control E15.5 embryos showing a multitude of hypertrophic GFP-electroporated (*green*) cells in BAF53a mutant brains that co-express the M-phase marker PH3 (*red*; *white arrow middle panel*). In the right panel, the number of yellow cells is dramatically reduced as nBAF expressing cells (green) exit the cell cycle and are no longer PH3 positive (red). *Scale bars, 25μm*.

To test the role of the BAF subunit switch *in vivo*, we used *in utero* electroporation to introduce vectors expressing the nBAF subunits BAF53b, BAF45b and CREST into BAF53a mutant and control cortices at E13.5. To evaluate the percentage of electroporated NPCs that exit the cell cycle (quit fraction), pregnant dams received an injection of BrdU at E14.5 (24 hours prior to sacrifice) and embryonic brains were analyzed at E15.5 (**Fig. 4d**). BAF53a mutant mice electroporated with a control GFP construct displayed a significantly reduced quit fraction compared to control mice, consistent with the observed cell cycle block following BAF53a deletion in NPCs. Electroporating BAF53a mutant mice with constructs expressing the nBAF subunits successfully rescued the quit fraction deficit (**Fig. 4e**), consistent with our observations *in vitro*. The majority of the nBAF-expressing mutant cells also no longer expressed the mitotic marker PH3 (yellow cells), further supporting the idea that nBAF expression rescues cell cycle block (specifically the arrest at G2/M) in BAF53a mutant mice and promotes cell cycle exit (**Fig. 4f**). In a series of control experiments, we also used *in utero* electroporation to overexpress nBAF and npBAF-specific subunits in wild type mice, in order to determine whether prolonged npBAF expression or premature nBAF expression was sufficient to alter the timing of cell cycle exit in the developing cortex (**Supplementary Fig. 6a**). We did not observe any significant differences in quit fraction with any of these manipulations as compared to GFP-only control electroporations (**Supplementary Fig. 6a, 6b**), suggesting that BAF53a repression is required to initiate differentiation and subunit switching. These striking results underline a causal role of BAF subunit switching in the regulation of cell cycle exit in the developing cortex. We show that BAF53a is required to maintain cell cycle progression and that its suppression leads to cell cycle arrest. Importantly, the nBAF subunit BAF53b is able to rescue the loss of BAF53a, releasing the cell cycle block by promoting cell cycle exit and subsequent acquisition of specific neural fates.

### BAF53a regulates cell cycle genes in the developing cortex

Chromatin remodelers such as the BAF complex regulate the epigenetic landscapes that establish cell type-specific gene expression patterns ^8^. To identify BAF53a-dependent target genes, we performed RNA-sequencing on E15.5 forebrains isolated from 4 control (*Nestin-Cre;Baf53a*^fl/+^) and 4 mutant (*Nestin-Cre;Baf53a*^fl/-^) mice. To determine significantly regulated genes, we applied a fold change cut-off of 1.5x and a significance threshold of FDR<0.1 (**Fig. 5a**). The sample-to-sample clustering and PCA analysis revealed that the datasets were highly reproducible, as they strongly clustered by genotype (**Fig. 5b, 5c**). Using these criteria, we identified 388 genes with increased expression and 200 genes with reduced expression in the BAF53a mutant brains (**Fig. 5d**). Using the DAVID software, we analyzed the 588 significantly regulated genes to associate Gene Ontology (GO) terms enriched in the dataset (**Fig. 5e**). Down-regulated genes were strongly enriched for GO terms associated with cell division, cell cycle and transcriptional regulation, whereas up-regulated genes were enriched for GO terms such as cell adhesion, ion transport, immune system process, axon guidance, cell migration and brain development, suggesting that deletion of *Baf53a* both suppresses the cell cycle and promotes neural differentiation. For example, key cell cycle genes such as *Ccng1, Ndc80, Ttk* and *Nusap1*, which regulate G2/M progression, were downregulated in mutant brains, as was the radial glial marker *Prom1*. Genes expressed in interneurons (e.g. *Dlx1, Dlx6*, and *Lhx6*), which migrate into the cortex from ventral telencephalic germinal zones, are also markedly reduced in expression upon *Baf53a* deletion, suggesting an impairment in the differentiation of those populations in BAF53a mutant mice. Furthermore, genes expressed in later-born excitatory neuronal populations and genes associated with glial differentiation, such as *Cux2, Pdgfra* and *Olig2*, were reduced in the absence of BAF53a. Immunostaining for glial lineages supported these findings, revealing an almost complete absence of OLIG2-expressing oligodendrocyte progenitors, as well as a depletion and displacement of BLBP and SOX9-expressing progenitors in mutant mice (**Supplementary Fig. 7a, 7b**). These data strongly support the phenotypic analysis we performed on the BAF53a mutant brains and suggest that BAF53a directly regulates genes involved in cell cycle regulation in the developing brain.

**Figure 5.**
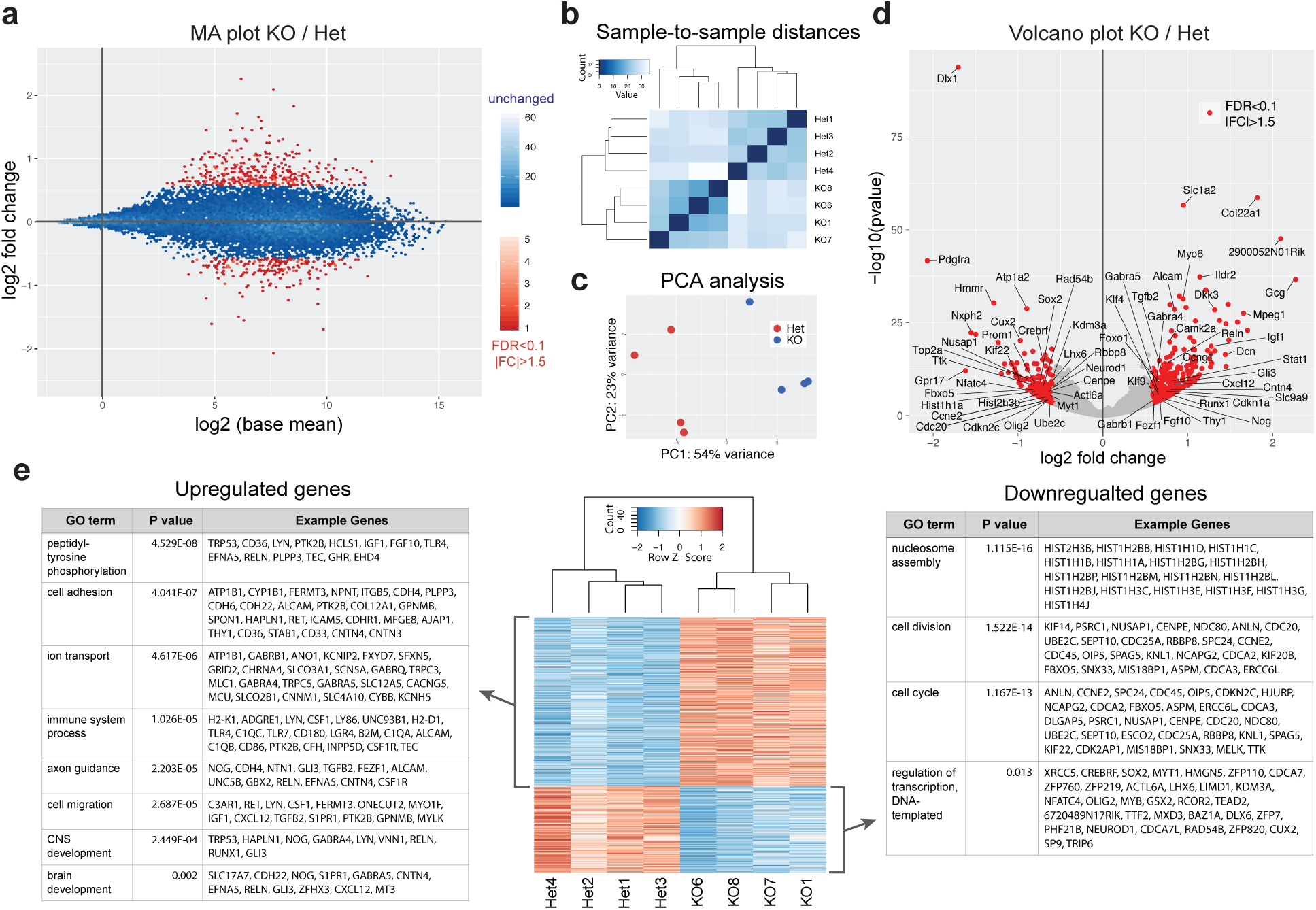
Loss of BAF53a leads to reduced expression of cell cycle regulators and enhanced expression of differentiation genes. **(a)** Genome-wide gene expression changes measured by RNA-seq in BAF53a mutant (*Nestin-Cre*;*Baf53a*^*fl/–*^, KO) and heterozygous control (*Nestin-Cre*;*Baf53a*^*fl/+*^, Het) forebrains at E15.5 (n=4). Magnitude and average (MA) plot (KO/Het) of RNA-seq data, displaying significantly changed genes in red (FDR <0.1 and fold change>1.5). **(b)** Sample-to-sample distance matrix reveals clustering of KO and KO replicates. **(c)** PCA plot highlighting reproducibility of biological replicates that cluster according to genotypes. **(d)** Volcano plot (KO/Het) with significantly upregulated and downregulated genes labelled in red. **(e)** Heat map of significantly upregulated and downregulated genes as well as Gene Ontology (GO) terms of biological processes associated with these genes.

### BAF53a regulates chromatin accessibility at neural enhancers

BAF chromatin remodeling complexes are thought to regulate gene expression by influencing chromatin accessibility at specific genomic loci ^37^. To determine the direct effects of *Baf53a* deletion on chromatin accessibility genome-wide, we performed ATAC-seq analysis on the forebrains of 4 control (*Nestin-Cre;Baf53a*^fl/+^) and 4 mutant (*Nestin-Cre;Baf53a*^fl/-^) mice. We found that 2521 accessible peaks were significantly changed (FDR > 0.1 and |fold change| > 1.5), representing 7% of the total number of accessible peaks across the genome in E15.5 forebrains (**Fig. 6a-d**). Using GREAT software to predict the function of these cis-regulatory regions, we associated significantly-regulated ATAC peaks to specific genes. GO terms analysis revealed a striking similarity with the results of our RNA-seq analysis (**Fig. 6e-f**). Lost accessibility peaks were associated with GO terms such as neuron fate commitment, stem cell maintenance, maintenance of cell number and regulation of Notch signaling. This finding is in line with our phenotypic analysis of NPCs in BAF53a mutant mice and reveals that BAF53a controls accessibility at discreet regulatory regions associated with genes that control NPC identity and numbers, such as *Pax6, Sox2, Ascl1, Notch1* and *Hes1* (**Fig. 6g**). We also found that accessibility was decreased at specific regulatory regions associated with gliogenesis genes (*Olig1/2, Nkx6-2*), interneuron genes (*Dlx1/2*) and upper-layer neuron genes (*Brn2*), consistent with our RNA sequencing data and immunostaining for markers of these cell types in Baf53a mutant brains. In contrast, gained accessibility peaks were associated with GO terms such as neuron development and differentiation, axon guidance and neuron projection guidance. This observation is in line with our phenotypic analysis of *Baf53a* mutant brains demonstrating premature induction of neuronal differentiation in mutant cortices at E15.5 (**Fig. 2e**). Increased NeuN staining in mutant brains is accompanied by increased accessibility at discreet regulatory regions associated with neuronal genes, including *Map2, Bdnf, Robo2* and *Dcc* (**Fig. 6h**). We also performed a chromatin feature analysis using ChromHMM software to characterize chromatin states at the ATAC peaks. We found that the majority of BAF53a-dependent ATAC peaks were at neural enhancers characterized by H3K4me1, H3K9ac and H3K27ac histone modifications in E15.5 forebrains (**Fig. 6f**). These results suggest that BAF53a-containing npBAF complexes directly influence accessibility to forebrain enhancers that regulate the expression of genes involved in the maintenance of NPC identity and the onset of neural differentiation.

**Figure 6.**
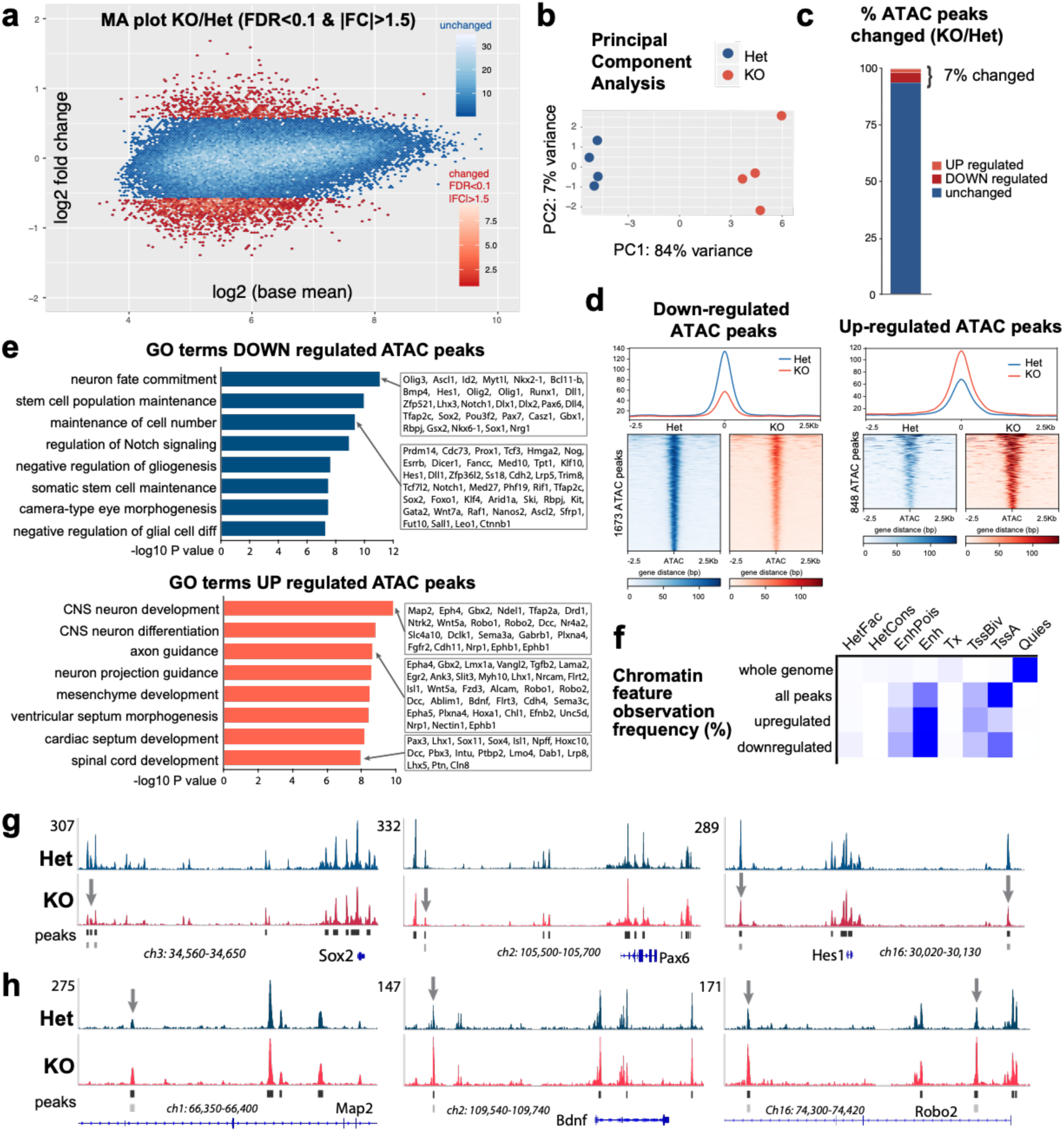
BAF53a regulates chromatin accessibility at NPC identity and neural differentiation genes. **(a)** Genome-wide chromatin accessibility changes as measured by ATAC-seq in BAF53a mutant (*Nestin-Cre*;*Baf53a*^*fl/–*^, KO) and heterozygous control (*Nestin-Cre*;*Baf53a*^*fl/+*^, Het) forebrains at E15.5 (n=4). Magnitude and average (MA) plot (KO/Het) of ATAC-seq data, displaying significantly changed peaks in red (FDR <0.1 and fold change>1.5). **(b)** PCA plot highlighting reproducibility of biological replicates that cluster according to genotypes. **(c)** Percentage of the total 39,041 ATAC peaks that are up or down regulated. **(d)** Genome-wide accessibility profiles at up- and down-regulated ATAC peaks in BAF53A KO and Het forebrains. **(e)** Analysis of Gene Ontology (GO) terms of biological processes associated with the genes that display gains or losses in accessibility in regulatory elements that control their expression. GREAT software was used to associate ATAC peak with a given gene. **(f)** Chromatin feature analysis reveals frequency of accessibility changes according to specific regulatory elements. The peaks which display changes in accessibility are most frequently associated with enhancer elements. ChromHMM was used to determine chromatin features using Encode data from E15 mouse forebrains. (**g, h**) Representative genome browser tracks showing changes in accessibility peaks in KO and Het forebrains at regulators of NPC identity and neural differentiation.

### BAF complexes regulate accessibility at neurogenesis transcription factor binding sites by opposing Polycomb-mediated repression

Changes in BAF53a-dependent accessibility were often found at discrete enhancer sequences rather than across all accessible regulatory regions associated with a given target gene (**Fig. 6g, 6h**). This suggested that accessibility at specific motifs may be regulated by the BAF complex in the developing forebrain. Motif analysis programs generally detect only the sequence motif for a transcription factor, but do not detect changes in occupancy induced by a genetic or environmental modification. However, a technique was recently developed that allows one to identify changes in transcription factor (TF) activity (BaGFoot), and we applied this method of analysis to the BAF53a target genes identified by ATAC-seq. Using BaGFoot analysis software to determine both changes in flanking accessibility, as well as changes in TF footprints, we found that members of the ASC, SOX, BRN and PAX families of TFs displayed reduced activity in BAF53a mutant brains (**Fig. 7a**). For example, ASCL1 motifs displayed reduced accessibility in BAF53a cKO forebrains compared to controls (**Supplementary Fig. 7c**). Furthermore, HOMER analysis revealed that ASCL1, BRN2, and SOX motifs were enriched in the peaks that displayed decreased accessibility upon loss of BAF53a (**Supplementary Fig. 7d)**. SOX2, strongly expressed in radial glial cells, is a key regulator of NPC maintenance and proliferation in the developing cortex ^38^. ASCL1 and BRN2 TFs also regulate progenitor proliferation and neuronal production during brain development ^39-41^. Moreover, the combination of MYT1L, ASCL1 and BRN2 can reprogram cellular identity and convert fibroblasts into neurons ^42^, further highlighting their central role in the regulation of neuronal differentiation. Interestingly, suppression of BAF53a in fibroblasts with miR-9/miR-124 reprograms human fibroblasts to neurons without a need for ASCL1 ^43^. Collectively, these data led us to propose that BAF53a is required to generate accessibility for neurogenic TFs to bind specific motifs and regulate target genes during forebrain development.

**Figure 7.**
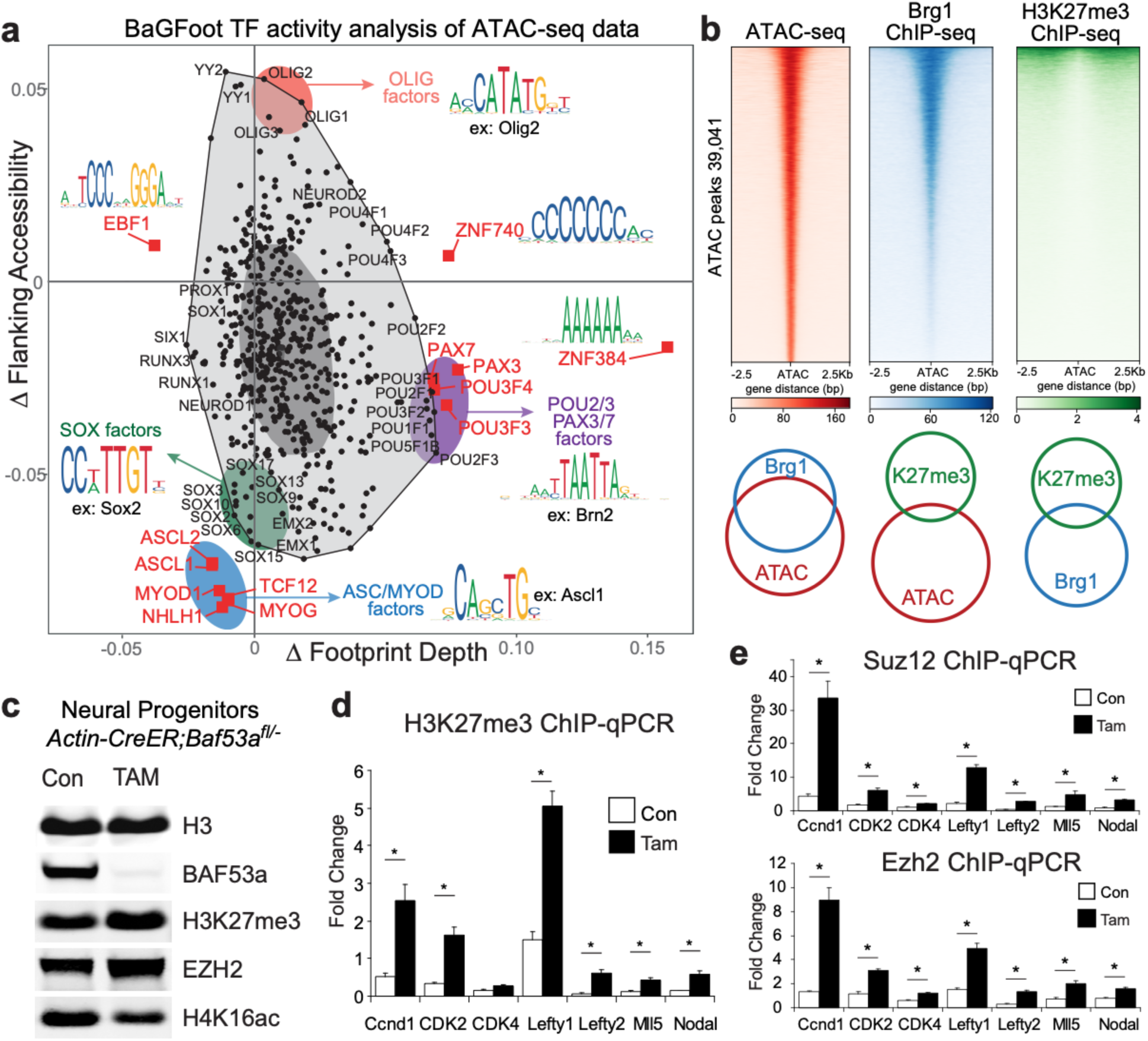
BAF53a regulates accessibility for neurogenesis TF activity through opposition to Polycomb repressive complex 2. **(a)** BaGFoot analysis of TF activity at BAF53a-dependent ATAC peaks. This analysis determines both changes in flanking accessibility, as well as changes in TF footprints. ASC, SOX, BRN and PAX families of TFs displayed reduced activity in BAF53a mutant brains. TF binding motifs for a given member of each family are displayed as examples. **(b)** Analysis of ChIP-seq and ATAC-seq datasets to characterize BRG1 and H3K27me3 levels across accessibility peaks in wild-type mouse forebrains. The majority of accessible regions are bound by BRG1, whereas very few accessible regions are marked with H3K27me3. Lower panel: overlap between accessible peaks, BRG1 binding and H3K27me3 marks. **(c)** Western blot analysis of H3K27me3 levels in *Actin-CreER*;*Baf53a*^*fl/–*^ NPCs treated with tamoxifen for 72 hours. **(d)** ChIP-qPCR analysis of H3K27me3 levels at cell cycle regulator genes, stem cell maintenance genes and *Hox* cluster genes. The levels of the H3K27me3 repressive marks are increased at cell cycle genes following *Baf53a* deletion in *Actin-CreER*;*Baf53a*^*fl/–*^ NPCs. n=3, error bars represent SEM, *p<0.05. **(e)** ChIP-qPCR analysis of SUZ12 and EZH2 levels at cell cycle regulator genes and stem cell maintenance genes. The levels of these Polycomb Repressive Complex 2 (PRC2) subunits are increased at cell cycle genes following *Baf53a* deletion in *Actin-CreER*;*Baf53a*^*fl/–*^ NPCs. The PRC2 complex places repressive H3K27me3 marks on histones. n=3, error bars represent SEM, *p<0.05.

Finally, we sought to determine the molecular mechanism through which BAF53a-containing BAF complexes regulate accessibility at these sites to control the expression of genes involved in maintaining NPC identity and cell cycle progression. In mouse embryonic stem cells, BAF has been shown to promote chromatin accessibility at enhancers and bivalent genes through eviction of the Polycomb repressive complex 2 (PRC2), which places the repressive H3K27me3 histone modification to repress gene expression ^44-46^. We analyzed BRG1 and H3K27me3 ChIP-seq data from wild-type forebrains and determined the overlap with the ATAC peaks measured in our datasets. We found that the majority of accessible peaks overlapped with peaks bound by BRG1, whereas very few accessible peaks were marked with H3K27me3 (**Fig. 7b**). This suggests that BAF complexes may generate accessible chromatin in the developing forebrain by restricting the placement of H3K27me3 by PRC2 complexes. To test this idea, we measured H3K27me3 levels in NPCs lacking BAF53a. We found that inducible BAF53a deletion led to increased global levels of H3K27me3 in cultured cells, as well as in the developing cortex at E15.5 (**Fig. 7c, Supplementary Fig. 7e**). Moreover, we performed H3K27me3 ChIP experiments in BAF53a mutant cells and found that H3K27me3 levels were increased at cell cycle genes (*Ccdn1, Cdk2* and *Cdk4*) (**Fig. 7d**). ChIP experiments for PRC2 complex subunits SUZ12 and EZH2 revealed similar increases at cell cycle genes and stem cell identity genes following BAF53a deletion (**Fig. 7e**). Taken together, these results suggest that BAF53a is required for npBAF complexes to directly oppose Polycomb at cell cycle genes and promote accessibility for TF binding (**Fig. 8**).

**Figure 8.**
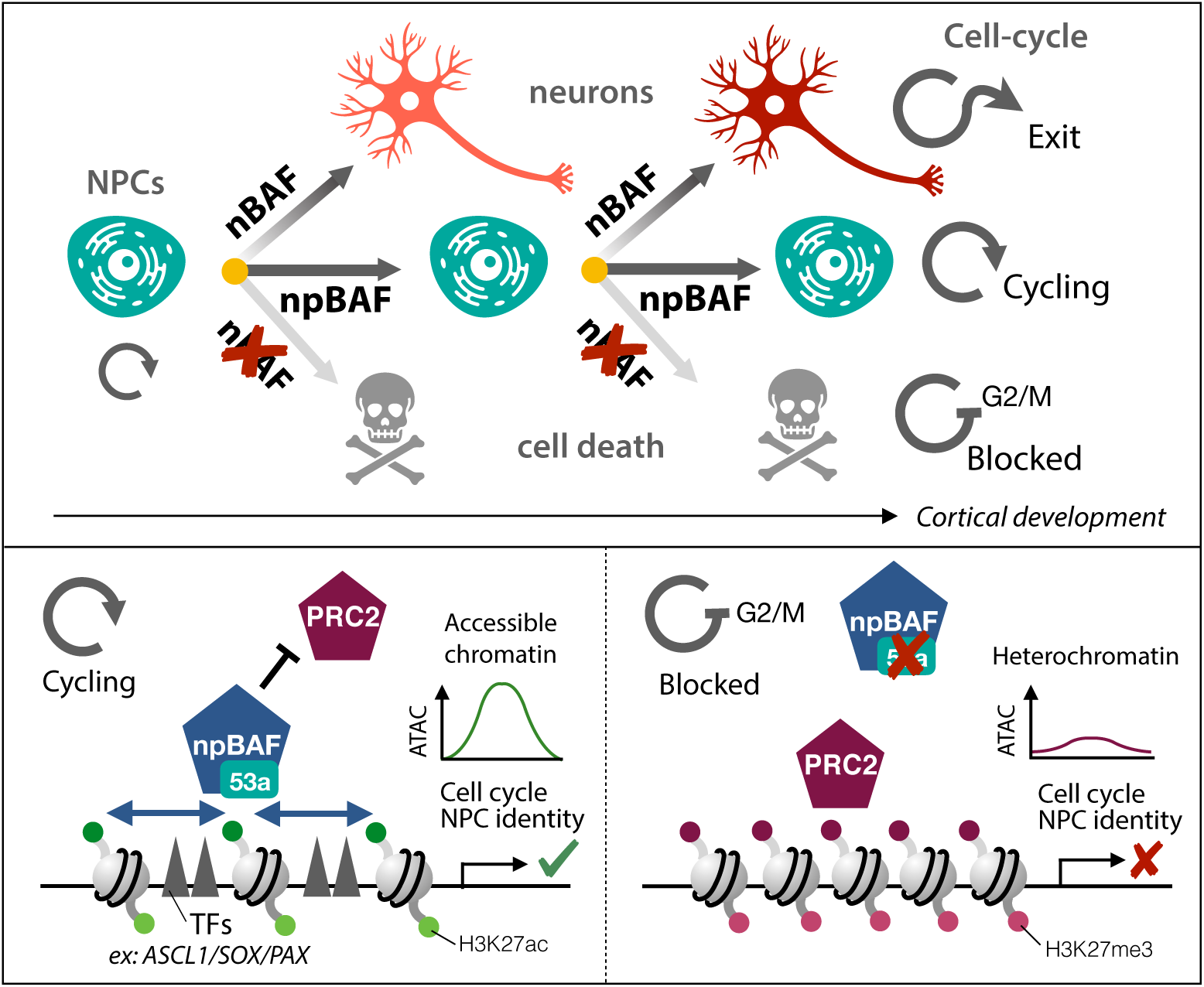
The npBAF to nBAF chromatin switch regulates cell cycle exit in the developing mammalian cortex. During cortical development, the switch from npBAF to nBAF chromatin regulators enables cell cycle exit and assures faithful execution of post mitotic neural differentiation programs in the cortex. NPCs that repress the npBAF subunit BAF53a, but fail to express the nBAF subunit BAF53b, remain blocked at the decatenation checkpoint and undergo apoptosis (upper panel). In cycling cells, continued expression of BAF53a-containing npBAF complexes generates accessibility at cell cycle and NPC identity genes through opposition to PRC2. When the npBAF subunit BAF53a is deleted, PRC2 places repressive H3K27me3 marks to repress the expression of cell cycle and NPC identity genes (lower panel).

## DISCUSSION

Our studies indicate that the npBAF to nBAF switch functions to promote the exit of neural progenitors from the cell cycle, acting as a checkpoint for the execution of neuronal differentiation programs during cortical development. The three key findings of our study are that 1) BAF53a-containing npBAF complexes directly regulate accessibility at genes required for normal cell cycle progression and NPC maintenance in the embryonic brain, 2) suppression of BAF53a is critical for progenitor cell cycle exit and differentiation onset, and 3) intrinsic and extrinsic mechanisms that normally regulate NPC proliferation cannot drive proliferation in the absence of BAF53a, whereas BAF53b, which normally replaces BAF53a, does overcome this cell cycle block (**Fig. 8**). In prior studies, we found that prolonging expression of BAF53a by deleting the miR-9/miR-124 binding sites in the 3’UTR of the *Baf53a* gene leads to increased progenitor proliferation ^6^. Here we showed that premature onset of miR-9/miR-124 expression *in vitro* phenocopies BAF53a loss, with both manipulations resulting in impaired cell cycle progression. Collectively, these observations lead us to propose that BAF53a-containing npBAF complexes are essential for NPC proliferation and that timely repression of BAF53a during normal neural development acts as a decisive signal to exit the cell cycle.

Proliferation in the developing nervous system involves the convergence of environmental cues and genetic programs balancing NPC maintenance with cell cycle exit. Despite extensive investigation, the precise mechanisms that allow NPCs to stop responding to pro-proliferative signals within their environment and instead exit the cell cycle have remained unclear. We sought to identify intrinsic molecular mechanisms that act to directly coordinate cell cycle exit in the developing cortex. Earlier studies suggested roles for npBAF complexes in NPC proliferation and set the stage for the hypothesis we tested here: that BAF subunit switching is an essential checkpoint that drives mitotic exit during corticogenesis. Brain-specific deletion of core BAF subunits BRG1 or BAF170 result in opposing phenotypes with respect to cortical thickness ^5,47^, suggesting that BAF complexes dynamically control brain size and neuronal content through the expression of specific subunits. npBAF complexes containing BAF53a have been proposed to play a general function in cell cycle maintenance in multiple cell types, as deletion of *Baf53a* leads to cell cycle exit in ES cells, hematopoietic stem cells, and keratinocytes ^9,Aizawa:2004jp; 48,49^. Complementary gain of function studies in the developing spinal cord have reported that extended expression of the npBAF subunits BAF45a or BAF53a is sufficient to promote proliferation, whereas expression of their homologs BAF45b and BAF53b, dedicated subunits of nBAF complexes, fail to do so ^5^. Taken together with our findings in this study, these data support the model that npBAF complexes play key roles in the maintenance of progenitor proliferation.

The cellular mechanisms by which npBAF complexes regulate proliferation have to date been poorly understood. Our genomic studies indicate that in the developing nervous system npBAF complexes containing BAF53a play a fundamental programmatic role in controlling cell cycle progression, by directly operating on genes that regulate both the G1/S transition as well as progression through G2/M. In line with these data, we found that deletion of *Baf53a* leads to cell cycle arrest at both G1/S and G2/M, characterized *in vivo* by a marked disorganization of the germinal zones accompanied by modest cell death and a dramatic increase in PH3+ cells throughout the VZ/SVZ. Upon BAF53a deletion, the expression of a number of genes involved in G2/M progression, including *Ccng1, Ndc80, Ttk, Nusap1*, and the haploid insufficient *Cenpe*, is coordinately altered ^50^. Owing to these coordinated alterations in the expression of mitotic checkpoint genes and the essential role of BAF complexes in topoisomerase II function ^31^, we propose that the increase in pH3+ cells we observe throughout the germinal zones of BAF53a mutant mice reflects a failure to pass the decatenation mitotic checkpoint. In other cell types, deletion of BAF subunits leads to G2/M arrest due to an inability of Topo-IIa to decatenate DNA prior to M-phase ^31^. Consistent with a mechanism of G2/M arrest in which Topo-IIa fails to bind DNA and in turn activates the decatenation checkpoint, we found that deletion of *Baf53a* results in a significant increase in anaphase bridges, a hallmark of failed Topo-IIa function reflecting improper segregation of daughter chromosomes. We also observed reduced expression of genes encoding members of the condensin complex in Baf53a mutants. Mutations in subunits of this complex have been previously associated with microcephaly and defects in decatenation at mitosis ^51^, further supporting the idea that BAF53a mutant NPCs are stuck at this checkpoint. Activation of the decatenation checkpoint arrests cells prior to the dephosphorylation of histone H3, thus explaining the accumulation of large pH3+ cells in the germinal zones of BAF53a mutant mice. Failure to pass this checkpoint may also disrupt interkinetic nuclear migration of dividing radial glial cells, whose nuclei normally descend to the ventricular surface during G2 to undergo mitosis. In line with this, we observed displacement of pH3+ and pVimentin+ radial glia away from the ventricle in BAF53a mutant brains, as well as decreased expression of the radial glial marker *Prom1*. Finally, if the decatenation checkpoint is not resolved, cells undergo apoptosis, which is highlighted by the increase in cleaved caspase 3 staining in BAF53a mutant brains.

A variety of intrinsic and extrinsic niche-dependent factors can promote proliferation of NPCs in the developing cortex. Some of the proteins transducing these signals, e.g. β-catenin, have also been shown to interact with BAF complexes ^52,53^. We reasoned that manipulating these factors in the context of *Baf53a* deletion would enable us to determine whether BAF53a acted downstream of known proliferative signals. In our study, none of the factors that we tested, including EGF, FGF, c-myc, SOX2, PAX6, and constitutively active β-catenin, could fully rescue the G1/S block upon *Baf53a* deletion, and all failed to rescue G2/M arrest. This observation suggests that repression of BAF53a and the resulting changes in gene expression, even in the presence of other proliferative signals, represents a molecular insurance policy allowing NPCs to reduce their responsiveness to mitogenic factors and faithfully exit the cell cycle following the G2/M transition. This idea is in line with previous studies suggesting that the responsiveness of intermediate progenitors to environmental cues is diminished after G2/M ^54,55^ and is supported by the upregulation of neural differentiation genes upon *Baf53a* deletion, including *Camk2a* and *Fezf1* ^56^, as well as the coordinated changes in chromatin accessibility revealed by our ATAC-seq analyses. *Baf53a* deletion results in decreases in accessibility at genes associated with maintaining progenitor identity (e.g. *Pax6, Hes1, Sox2*) and concomitant increases in accessibility at genes associated with neuronal differentiation (e.g. *Map2, Ptbp2*) and axon development (e.g. *Wnt5a, Nrcam*). Moreover, in the ATAC peaks displaying decreased accessibility upon BAF53a deletion, we found an enrichment in TF motifs associated with NPC maintenance and proliferation, including SOX2 and ASCL1 ^38,39^. These data indicate that BAF53a deletion may set differentiation programs in motion and they are reinforced by the increased MAP2ab staining we observe upon tamoxifen administration in our *in vitro* experiments, as well as the increased NeuN staining observed in BAF53a mutant cortices *in vivo*. Importantly, both *in vitro* and *in vivo*, expression of BAF53b alone or in combination with other nBAF subunits is sufficient to rescue cell cycle block, promoting exit from the cell cycle and differentiation. This exciting observation supports the model that loss of BAF53a induces cell cycle exit, and when replaced by the neuron-specific BAF53b, neural differentiation programs are consolidated and allowed to progress. Expression of nBAF subunits in wild-type mice does not alter cell cycle exit, implying that repression of BAF53a is required to initiate subunit switching and differentiation, but premature inactivation of BAF53a leads to a permanent block in cell cycle progression. These findings are also consistent with our earlier studies showing that post-mitotic nBAF complexes containing BAF53b do not regulate cell cycle genes, instead targeting genes involved in the regulation of neuronal function ^4,27^. The failure to exit the cell cycle seen in BAF53a mutants also leads to a loss of later born glial cells, which is supported by previous observations highlighting the role of the BAF complex in targeting Olig2 TFs to initiate oligodendrocyte differentiation ^57^.

We have shown that the BAF complex regulates cell cycle progression in the developing brain by shaping chromatin accessibility at genes controlling NPC proliferation and differentiation. The question remains, however: how do BAF53a-containing npBAF complexes orchestrate these coordinated gene expression changes to promote progenitor proliferation? In BAF53a mutant cortices, we observed a significant increase in H3K27me3, a repressive chromatin mark deposited by PRC2. This observation, coupled with our ChIP-qPCR data showing increased H2K27me3 and PRC2 binding at cell cycle genes in BAF53a mutant cells, suggests that BAF53a regulates cell cycle progression and NPC maintenance in part by antagonizing PRC2. The proliferation of progenitors is thus regulated by a direct opposition between npBAF complexes and PRC2 at BAF53a target sites. Our findings are consistent with previous studies in ES cells and fibroblasts demonstrating that deletion of *Brg1* results in aberrant H3K27me3 near BRG1 peaks, and that recruitment of BAF complexes to repressive chromatin leads to PRC2 eviction and loss of H3K27me3 marks within minutes ^44,45,58^. In the developing cortex, it was recently shown that disruption of PRC2 via conditional deletion of the core subunit *Eed* reduces H3K27me3 levels, altering the time course of NPC differentiation and resulting in the precocious generation of later-born upper layer neurons ^29^. In line with these findings, we observe the opposite phenotype in our BAF53a mutants, where cell cycle progression is blocked and the generation of later-born cell types, including upper layer neurons, is broadly impaired. Taken together, our findings highlight a central role for chromatin regulators in the output of NPCs during brain development. We propose that antagonism between the BAF and Polycomb complexes controls cell cycle exit by altering the balance of repressive and active chromatin states at cell cycle genes.

Exome sequencing studies have identified mutations in BAF complex subunits in patients with various neurodevelopmental disorders, including ASDs, selective speech disorders, and primary microcephaly syndromes characterized by intellectual disability, underscoring key roles played by these complexes in neural development ^12,14,16,59-62^. Indeed, recent studies have found that monoallelic mutations of *ARID1B* are the most frequent cause of de novo intellectual disability ^63^. Our data here and in earlier publications indicates that Polycomb opposition, in the form of direct ATP-dependent eviction of PRC1/2, is a key function of BAF complexes that offers an novel point for therapeutic intervention ^44,45^. The nucleosome remodeling activity of SWI/SNF, on the other hand, is redundant with other chromatin remodeling complexes in both yeast and mammals ^64^, and is thus a less likely mechanism to explain the developmental outcomes of BAF mutations. Future biochemical studies defining the precise steps in the ATP-dependent direct eviction of Polycomb may lead to the recognition of new therapeutic avenues for common neurodevelopmental disorders resulting from BAF subunit dysfunction or mutation.

## METHODS

### Animals

All animal experiments were approved by the Stanford University and UCSF Institutional Animal Care and Use Committees and conducted in accordance with NIH guidelines for the care and use of laboratory animals. Heterozygous mice carrying a germline deletion were intercrossed to generate BAF53a null mice. *Baf53a*^fl/fl^ mice ^49^ were mated with *Baf53a*^*+/-*^*;Nestin-Cre* mice ^65^ to generate conditional knockout pups. Transgenic mice expressing constitutively active β-catenin ^22^ were bred with *Nestin-Cre* mice. All the lines have been backcrossed to C57BL/6J background for a minimum of 7 generations. Genotyping was performed by PCR of tail DNA at three weeks when weaning.

### *In utero* electroporation

Timed-pregnant Swiss Webster dams (Charles River) or dams carrying *Baf53a* mutant alleles (*Baf53a*^*fl*^ *or Baf53a*^*null*^) at E13.5 were deeply anaesthetized using isofluorane inhalation anesthesia, and the uterine horns were exposed. 1 μl (concentration 3μg/μl) of control plasmid (GFP), or a mix of GFP and a polycistronic nBAF (pCAG-53b-CREST-45b) or npBAF (pCAG-53a-SS18-45d) vector was injected into the lateral ventricles using a sterile glass microinjection pipette. The E13.5 embryos were then electroporated through the uterine wall using square pulse electroporator (BTX 830) with five 50 msec pulses of 38V. Following electroporation, the uterine horns were placed back into the dam for the embryos were to recover and grow. For quit fraction analyses, the pregnant dam was injected intraperitoneally with 50mg/kg BrdU at E14.5 (24h post-electroporation), and embryonic brains were harvested 24 hours post-injection at E15.5. A minimum of n=3 embryos from at least two different litters were used for each analysis.

### Neural progenitor cultures

Single cells were dissociated from E12.5 embryonic forebrains and cultured in suspension in DMEM/F-12 (Life Technologies, 10565-042) containing N2 supplement (Life Technologies, 17502-048), B27 supplement (Vitamin A minus, Life Technologies, 12587-010), HEPES (Invitrogen, 15630080), non-essential amino acids (Invitrogen, 11140050), GlutaMAX (Invitrogen, 35050061), Penicillin/Streptomycin (Invitrogen, 15140122) and ß-mercaptoethanol (Invitrogen, 21985023). bFGF (Invitrogen, 13256-029) and EGF (Invitrogen, PHG0311) growth factors were used as mitogens to maintain proliferative NPC cultures. For passaging the neurospheres were digested with papain (Worthington, LS003126) and mechanically dissociated into single cells. NPCs were subsequently grown on poly-ornithine coated dishes for adherence. For the in vitro rescue experiments, adherent cultures of *Actin-CreER*;*Baf53a*^*fl/–*^ NPCs were infected with lentiviruses and selected with puromycin for 3 days followed by 3 days of tamoxifen (TAM) treatment to delete *Baf53a*. Cell were maintained in proliferative conditions by supplementing the culture media with EGF and bFGF growth factors.

### Immunohistochemistry

For immunocytochemistry, cells were fixed with 4% PFA for 10 min at 4°C. Antibodies used were Nestin (DSHB; Rat-401), Map2ab (Millipore, MAB378Tuj1), BAF53a (Novus, NB100-61628), BAF53b (Crabtree lab). Mouse embryonic brain tissue was collected at various time points from electroporated wild-type and BAF53a mutants animals, fixed in ice-cold 4% PFA (EMS, 15710) for 30 min on ice (or overnight at 4°C for E18-P0 brains), and cryoprotected in 30% sucrose before embedding in OCT (Sakura, 4583) and sectioned on a Leica cryostat at 16 *μ*m. Sections were kept frozen at −80°C until needed for staining. Sections were blocked in 5% serum in 0.25% TritonX/PBS and incubated in primary antibody overnight. Primary antibody binding was detected using Alexa fluorophore-conjugated secondary antibodies (ThermoFisher Scientific). For BrdU immunostaining, brain sections were first stained for all other antigens and then post-fixed in ice-cold 4% PFA before undergoing antigen retrieval (30min incubation in 2N HCl at 37°C) and then incubated in anti-BrdU primary antibody overnight. Brain sections were incubated for 30 min in 95°C citrate buffer (pH=6) prior to staining for CUX1. Antibodies used for immunofluorescent tissue staining include KI67 (BD Pharmingen, 550609), PH3 (Millipore Sigma, 160189), BrdU (Abcam, ab6326), CC3 (Cell Signaling Technology, 9664), GFP (Abcam, ab13970), pVim (Abcam, ab22651), SOX2 (Santa Cruz, sc17320), Tbr2/Eomes (Thermo Fisher, 14-4875-82), DCX (Santa Cruz, sc8066), MAP2ab (Millipore Sigma, MAB378), NeuN (Millipore Sigma, MAB1281), TBR1 (Abcam, ab31940), CTIP2 (Abcam, ab18465), SATB2 (Abcam, ab51502), CUX1 (Santa Cruz, sc13024), BRN1 (Novus Biologicals, NBP1-49872), SOX9 (R&D Systems, AF3075), BLBP (Abcam, ab32423), OLIG2 (Millipore Sigma, AB9610).

### Microscopy and Image Analysis

Images were acquired on an upright epifluorescent Zeiss Axioscope Imager.M2 using 20x objective, as well as a Keyence BZ-X700 microscope, and processed using Adobe Photoshop CC2019. For quantitative analysis, cells within the 20x field view were counted manually using ImageJ (Public Domain, BSD-2) Cell Counter application. Cell counts were performed on a minimum of 3 region-matched brain sections per animal. Results were plotted and analyzed using Prism 8 (GraphPad).

### FACS analysis

NPC cultures were labeled with BrdU (BD Pharmingen, 559616) for 1 hour or 2 hours, respectively. Trypsinized cells were processed for BrdU-FITC and 7AAD (7-Aminoactinomycin D) staining following the manufacturer’s protocol (BD Pharmigen, 559616). FACS was run on a Scanford analyzer (Stanford FACS Core Facility) and data was analyzed using Flowjo software.

### Immunoprecipitation

Immunoprecipitations were performed on 250ug of nuclear extracts prepared as previously described (Kadoch et al. 2013). 2.5 micrograms of immunoprecipitation antibodies were incubated with nuclear extracts rotating at 4 degrees overnight, and then protein G Dynabeads were used to isolate Ig-protein complexes with 2h incubation rotating at 4 degrees.

### Western blot

The following antibodies were used for the western blots: mouse anti-Brg1 (Santa Cruz, sc-17796, 1:1000), rabbit anti-Suz12 (Cell Signaling, 3737, 1:1,000), rabbit anti-actin (Sigma, A2066, 1:500), rabbit anti-H3 (Abcam, 1:2,000), rabbit anti-BAF53a (Novus, NB100-61628, 1:1,000), rabbit anti-Epc-1 (Abcam, ab5514, 1:1000), rabbit anti-Tip60 (Millipore, 07-038, 1:500), mouse anti-β-actin (Abcam, ab6276, 1:1000). Antibodies against BAF subunits were generated in our laboratory and used at the following concentrations: BAF53b (1:500), BAF45d (1:2,000) and BAF155 (1:1,000). The secondary antibodies were goat anti-rabbit or mouse conjugated with IRDye 680RD or 800CW. Images were taken and quantified by Odyssey Quantitative Fluorescence Imaging Systems.

### Differential salt extraction of chromatin-associated proteins

Nuclei from NPCs were isolated using Buffer A (25 mM HEPES, pH 7.6, 5 mM MgCl_2_, 25 mM KCl, 0.05 mM EDTA, 10% glycerol, 0.1% NP-40) and aliquoted into separate tubes for further treatment. Pellets were resuspended in Buffer A with different NaCl concentrations as indicated and were rotated at 4°C for 30 min, followed by high-speed centrifugation for 15 min to pellet insoluble chromatin. Nuclear lysates were removed and mixed with 4X gel loading dye for western blot analysis. Chromatin fraction was solubilized for western blot by sonication.

### Anaphase bridge analysis

Analysis was performed as previously described (Dykhuizen et al., 2013). 72h after Tamoxifen treatment, cells were fixed with 4% para-formaldehyde for 20 min, washed with PBS and DNA content was visualized by staining with 49,6-diamidino-2-phenylindole (DAPI 1µg/ml; Sigma) to detect and count anaphase bridges.

### ATPase assay

Assay was performed as previously described (Dykhuizen et al., 2013). Immunoprecipitates were washed with 10mM Tris-HCl, pH7.5, 50 mM NaCl, 5 mM MgCl_2_, 1 mM DTT and re-suspended in assay buffer (10 mM Tris-HCl, pH 7.5, 50 mM NaCl, 5 mM MgCl_2_, 20% glycerol, 1 mg/ml BSA, 0.2 mM ATP, 20 nM plasmid DNA, 50 mCi ml^-1^ g-^32^P ATP, 1 mM DTT, protease inhibitors). The reaction was agitated at 37°C for 1 h. Reaction mixtures were spotted onto PEI cellulose plates at the indicated timepoints and thin layer chromatography was performed in 0.5 M LiCl and 1 M formic acid. The plates were dried and imaged using phosphorimaging.

### Glycerol gradient

The glycerol gradient was prepared in HEMK buffer (25mM HEPES, 0.1mM EDTA, 12.5mM MgCl_2_, 100mM KCl) using a Hoefer SG 15 gradient maker (GE Healthcare Bio-Sciences). 300µg of nuclear extract from *Baf53a*^f/-^:Actin-CreER cells +/-tamoxifen was loaded onto the top of the gradient and separated by ultracentrifugation at 40,000 rpm for 16 hours at 4°C. 500µl of fractions were collected from top to bottom and precipitated by TCA. Precipitates were washed 2 times with acetone, air-dried and resuspended in 1XLDS loading buffer for western blotting.

### Plasmid construction and viral preparation

cDNAs of BAF53a, BAF53b and all other genes used in this study were cloned downstream of the EF1a promoter in a lentiviral construct with puromycin, hygromycin or blasticidin selection. The miR-9/-124 lenti-viral construct was reported before (Yoo et al., 2011), Lenti-X 293T cells (Clontech, 632180) were transfected with appropriate amounts of lenti-viral constructs, psPAX2 and pMD2.G using PEI (Polysciences, 24765-2) after reaching 70-80% confluence. Medium was replaced 16 hours after transfection and virus was harvested after 48 hours. Virus was filtered and spun down by ultracentrifuge for enrichment. Pellet was resuspended for infection. For in utero electroporation experiments, polycistronic vectors were cloned to generate pCAG-53b-CREST-45b (nBAF) and pCAG-53a-SS18-45d (npBAF) constructs which were co-electroporated with reporter pCAG-GFP plasmids.

### RNA-seq and data analysis

RNA-seq reads were processed by mapping to the mm10 reference mouse genome using Kallisto, Differential peak calls were made using DESeq2^45^ and the requirement was set at fold changes of >1.5-fold in either direction and FDR-corrected *P* < 0.10. DAVID software was used to determine Gene Ontology (GO) terms for biological functions associated with the up- & down-regulated genes.

### ATAC-seq and data analysis

ATAC-seq reads were processed by mapping to the mm10 reference mouse genome using Bowtie2, rejecting reads that contained more than a single mismatch. Duplicate reads were discarded, leaving only unique reads. Peak calling was performed by MACS2. Differential peak calls were made using DESeq2 and the requirement was set at fold changes of >1.5-fold in either direction and FDR-corrected *P* < 0.10. RPM values in genome tracks are the mean values across replicates from each condition. Deeptools was used to produce heatmaps of mean read density across all peaks. ChromHMM was used for feature analysis using 12 states based on previously published mouse E15.5 forebrain ChIPseq data obtained from Encode. GREAT software was used to annotate ATAC peaks with genes to determine Gene Ontology (GO) terms associated with changes in accessibility.

### BaGFoot Footprint depth and Flanking accessibility calculation

We used TF binding motifs from the JASPAR motif database (Khan et al., 2018), 579 motifs in total. TF footprint depth (FPD) was defined as:

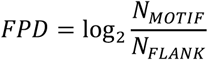

where *N*_*MOTIF*_ is the average number of Tn5 inserts in [-5 bp, +5 bp] window around motif center and *N*_*FLANK*_ is the average number of Tn5 inserts in the [-55 bp, −6 bp] U [6 bp, 55 bp] window around the motif center, calculated using motifs found in differentially accessed peaks. Flanking accessibility (FA) was defined as:

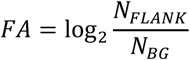

where *N*_*BG*_ is the number of Tn5 inserts in the background region defined as an interval [-200 bp, −250 bp] U [200 bp, 250 bp] around the motif center. Only motifs found within ATAC-seq peaks were used in the calculations. To visualize changes in FPD and FA between HET and KO we used bagplot ^66^.

### Homer motif analysis

We performed motif enrichment analysis using HOMER v4.11^67^. We constructed genomic intervals [–250 bp, 250 bp] around summits of peaks exhibiting significant change in accessibility between KO and HET samples. These intervals were used as input for findMotifsGenome.pl that was run with “-size given” option and other default parameters. We performed motif enrichment analysis for all peaks and separately for peaks exhibiting decreased and increased accessibility in KO relatively to WT.

### ChIP qPCR

ChIP experiments were performed as previously described (Hathaway et al., 2013). For RT-qPCR, DNA samples were prepared using the SensiFAST SYBR Lo-Rox kit (Bioline, BIO-94020), according to the manufacturer’s instructions. Analysis of qPCR samples was performed on a QuantStudio 6 Flex system (Life Technologies).

### Statistics

All data are presented as mean ± SEM. Pairwise comparisons were analyzed using a two-tailed Student’s t test. P values of less than or equal to 0.05, were considered statistically significant.

## Data availability

RNA and ATAC sequencing datasets of E15.5 forebrains generated and analyzed in this study will be made available in the Gene Expression Omnibus (GEO) repository. Published sequencing datasets were obtained from the Encode project (Reference epigenome in mouse forebrain at E15.5: ENCSR438IXZ). Other data and materials are available from the authors upon request.

## ACKNOWLEDGMENTS

We thank all members of the Crabtree and Panagiotakos labs for helpful discussions as well as S. Jessberger and K. Majzoub for helpful comments on the manuscript. Microscopy was performed using the Stanford Cell Sciences Imaging Facility. Special thanks to the Villeda lab at UCSF for sharing their imaging equipment. Next generation sequencing was performed using the Stanford Functional Genomics Facility. FACS analysis was performed using the Stanford Shared FACS Facility. This work was supported by an NIH grant NS046789 (G.R.C), funding from the Howard Hughes Medical Institute (G.R.C.), and generous support by the UCSF Program for Breakthrough Biomedical Research, Sandler Foundation (G.P.). G.R.C. is an HHMI Investigator. S.M.G.B. is supported by the Sir James Black GSK fellowship. G.P. is funded by the UCSF Sandler Faculty Fellows Program, and the Larry L. Hillblom Foundation provided support to R.P.

## AUTHOR CONTRIBUTIONS

G.P. and G.R.C. supervised all aspects of these studies. S.M.G.B., R.P., J.T., G.R.C., and G.P. designed and analysed the experiments. Experiments were executed by S.M.G.B., R.P., and J.T. with support by A.K., Y.T., and E.M. The sequencing data was analyzed by S.M.G.B. and A.K.; S.M.G.B., R.P., J. T., G.R.C., and G.P. wrote the paper with help and feedback from the rest of the co-authors.

