## Supplementary Figures for "The npBAF to nBAF Chromatin Switch Regulates Cell Cycle Exit in the Developing Mammalian Cortex"

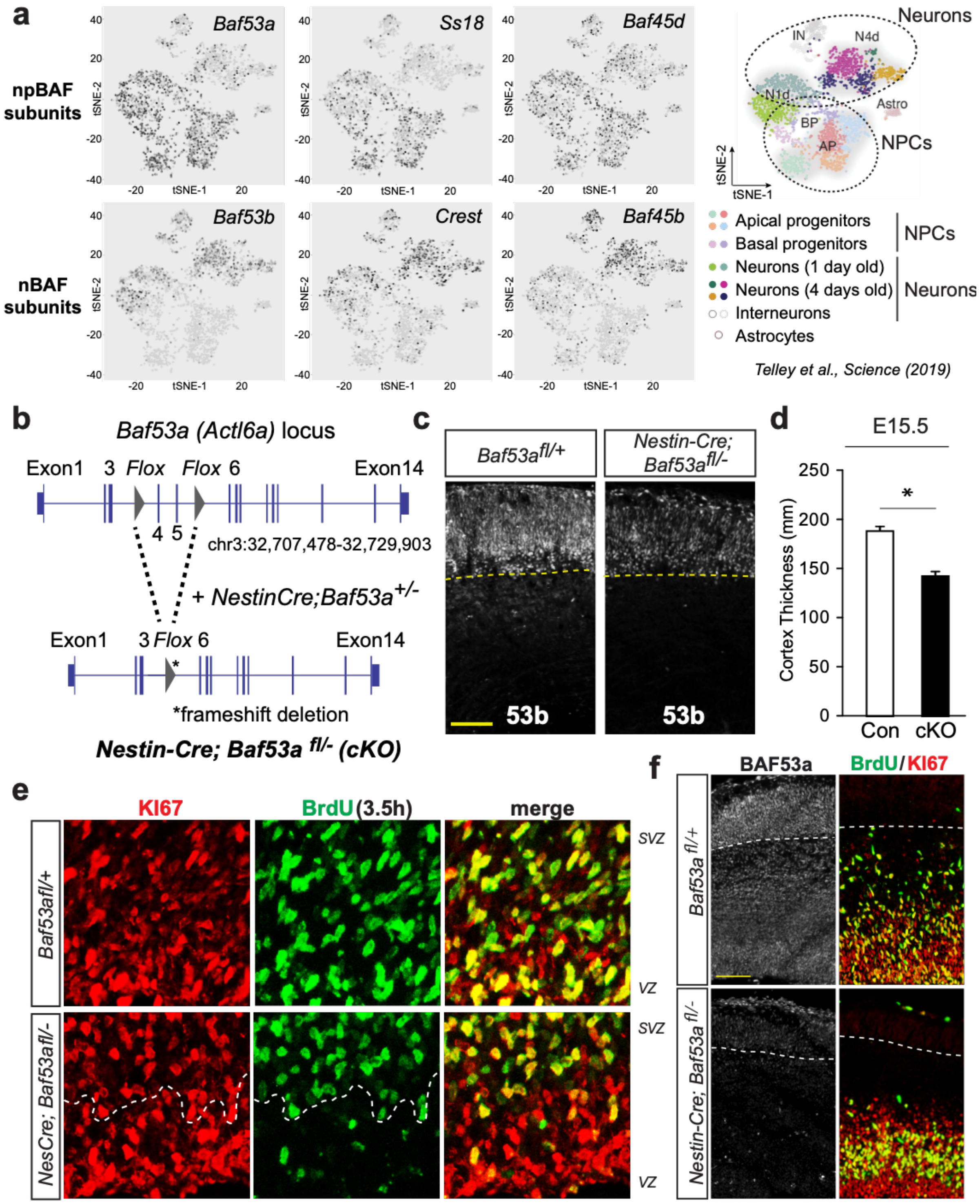


**Supp. Figure 1 (related to Figure 1). BAF53a is required for normal neural development.**
(**a**) Single cell RNA-seq data from Telley et al. Science (2019). Expression pattern of npBAF and nBAF subunits during embryonic neurogenesis from E12-E15.

(**b**) Schematic representation of the BAF53a (*Actl6a*) locus after incorporation of *loxP* (*flox*) sequences flanking exons 4 and 5 (Krasteva et al., 2012). By breeding *Baf53a*$^{fl/-}$ mice with *Nestin-Cre* expressing mice we generated brain specific BAF53a mutant mice.
(**c**) Immunofluorescence staining of coronal sections through E15.5 mouse cortex showing robust BAF53b expression in the cortical plate of control and BAF53a mutant brains. *Scale bar, 50µm*.
(**d**) Quantification of cortical thickness in E15.5 control and BAF53a mutant cortices. n=3, error bars represent SEM, *$p<0.05$.
(**e**) Immunostaining of E14.5 cortex for the proliferative marker KI67 (*red*) and short-term BrdU labeling (*green*, 3.5h pulse) highlighting a lack of marker co-localization in the ventricular zone of BAF53a mutants. *Scale bar, 50µm*.
(**f**) Immunostaining of E15.5 cortex for the proliferative marker KI67 (*red*) and short-term BrdU labeling (*green*, 3.5h pulse). *Scale bar, 50µm*.

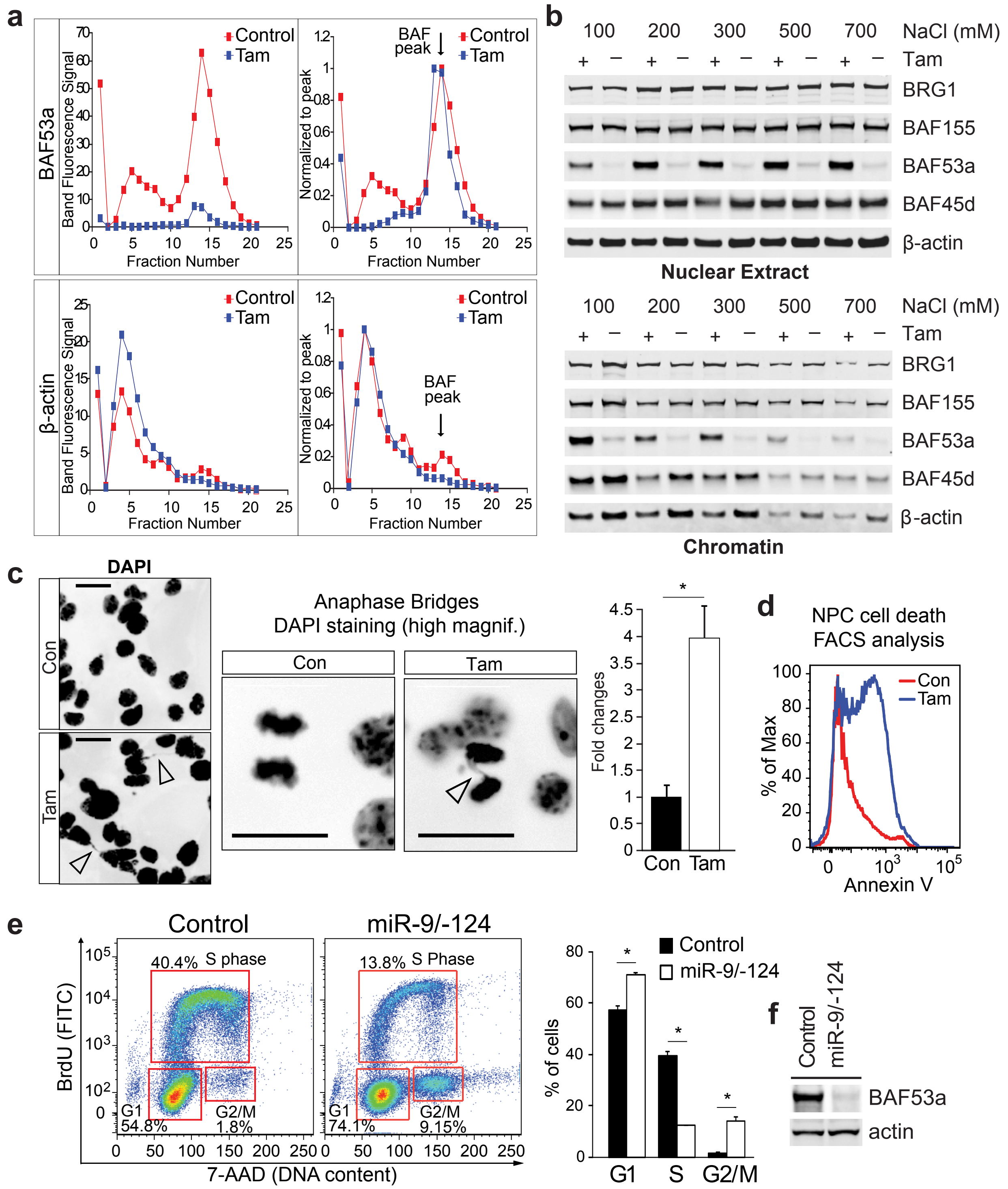


**Supp. Figure 2. Deletion of *Baf53a* does not affect BAF complex formation but results in an increase in anaphase bridges.**

(**a**) Quantification of BAF53a and β-actin signals from glycerol gradient immunoblots shown in **Figure 1d** from control and BAF53a mutant cells. Signals were quantified and normalized to the peak value. Protein signal peaks from fraction 13-15 correspond to the BAF complex (*down arrows*).

(**b**) Western blots examining the chromatin association of BAF complexes. Nuclei were prepared by hypotonic lysis and nuclear proteins were dissociated using varying concentrations of NaCl. Nuclear extract (supernatant) and chromatin (pellet) fractions were collected separately and analyzed by immunoblot using antibodies to different BAF complex components. **+**, control cells; **–**, BAF53a mutant cells.
(**c**) DAPI staining showing the decatenation defect following *Baf53a* deletion in neurospheres. Arrowheads indicate the DNA bridges (*left panel*) and M phase bridges (*right panel*), which are more often observed in the BAF53a mutant cultures. *Scale bar, 20μm*. Quantification of the relative number of DNA bridges between control and BAF53a mutant cells. n=3, error bars represent SEM, *p<0.05.
(**d**) FACS analysis for cell death using Annexin-V staining of NPCs following Baf53a deletion.
(**e**) Cell cycle FACS analysis of untransduced *Baf53a*$^{f/+}$ neurospheres and *Baf53*$^{f/+}$ neurospheres in which miR-9/-124 were prematurely expressed. Cells were incubated in BrdU 2 hours prior to collection and the fraction of cells in each stage of the cell cycle was assessed. Right panel: Quantification of the experiment. n=3, error bars represent SEM, *p<0.05.
(**f**) Western blot showing reduced levels of BAF53a in *Baf53a*$^{f/+}$ neurospheres expressing miR-9/-124.

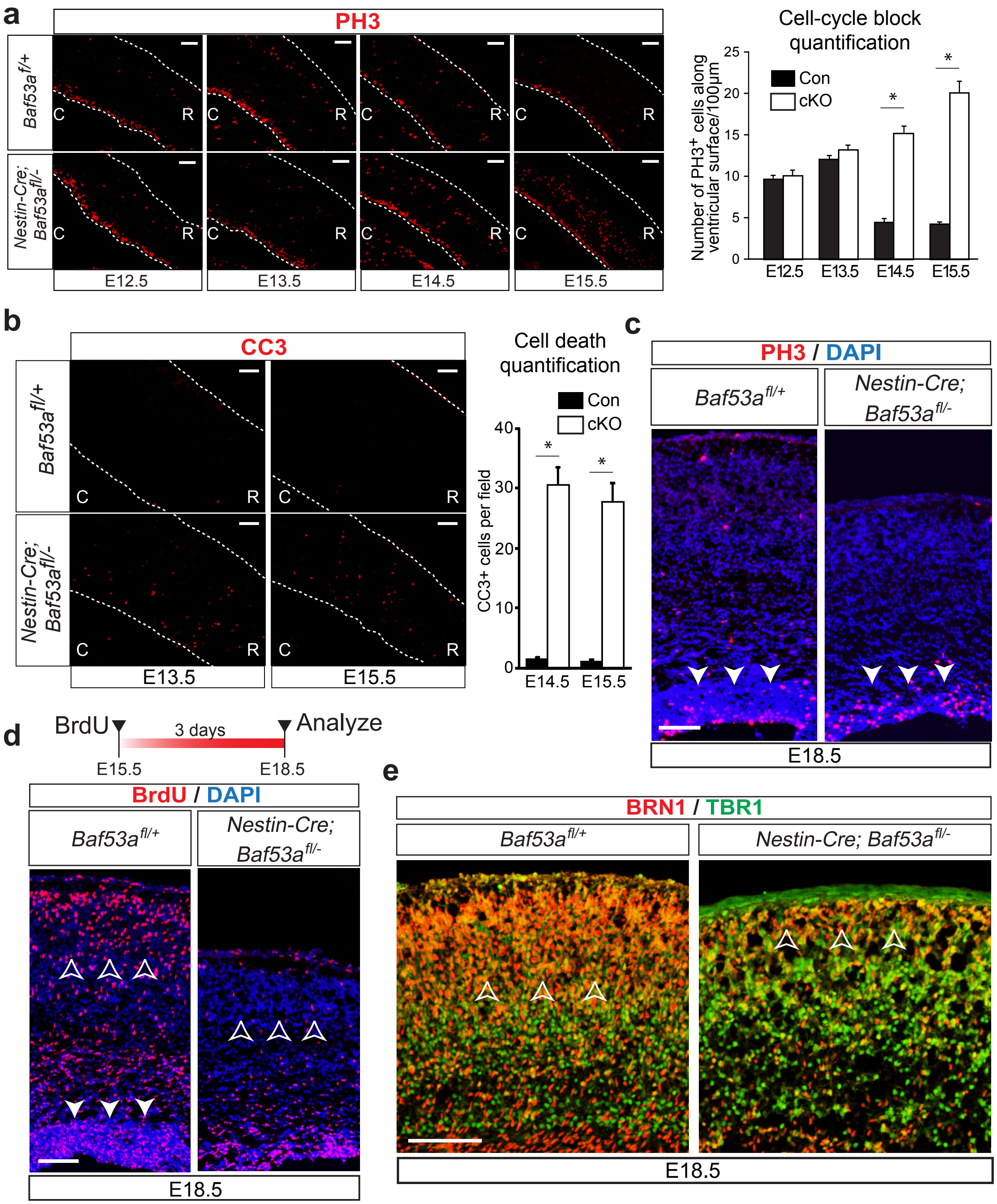


**Supp. Figure 3 (related to Figure 2). Deletion of *Baf53a* leads to elevated pH3 staining and increased apoptosis.**

(**a**) PH3 staining (*red*) of sagittal cortical sections from E12.5 to E15.5 BAF53a mutant and control mice. Dotted line contours the cortex spanning the ventricular wall to the pial surface. ***R**, Rostral; **C**, Caudal. Scale bar, 50µm*. Right panel: Quantification of PH3 (S10P, *red*) positive cells along the ventricular surface from E12.5 to E15.5. n=3, error bars represent SEM, *p<0.05.

(**b**) Cleaved Caspase 3 (*red*) staining of E13.5 and 15.5 sagittal brain sections from *Nestin-Cre; Baf53a*$^{f/-}$ and *Baf53a*$^{f/+}$ embryos. *Scale bar, 100μm.* Right panel: quantification of CC3+ cell number in control and cKO brains. n=3, error bars represent SEM, *p<0.05.
(**c**) Immunofluorescence staining of coronal sections through E18.5 mouse cortex showing persistent expression of the M-phase marker PH3 in BAF53a mutant brains. *Scale bar, 50μm*.
(**d**) Immunostaining for BrdU+ (*red*) expressing cells in BAF53a mutant and control brains. The thymidine analog was administered at E15.5 and tissue harvesting was performed at E18.5 to determine the fate of the proliferating NPCs labelled at E15.5. *Scale bar, 50μm*.
(**e**) Immunostaining for BRN1 (*red*) and TBR1 (*green*) expressing cells in BAF53a mutant and control brains. BRN1-expressing cells are late-generated neuronal cell types which are absent from BAF53a mutant brains.

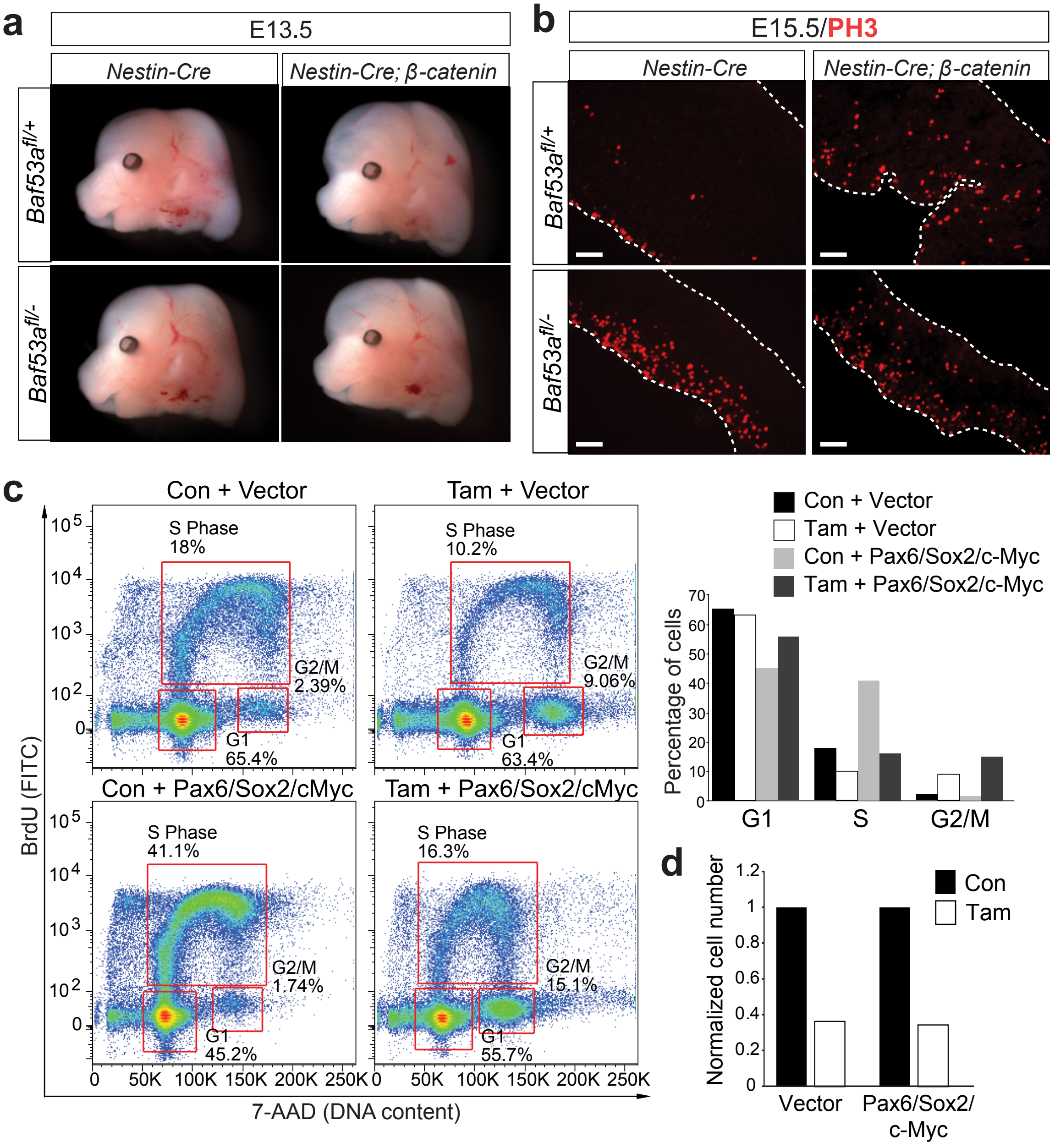


**Supp. Figure 4 (related to Figure 3). Activation of β-catenin does not rescue deletion of BAF53a**

(**a**) Microscopic images of E13.5 embryonic heads from Baf53a mutant and controls with or without the β-catenin transgene.

(**b**) Photomicrographs of sagittal sections from Baf53a mutant and control mice with or without the β-catenin transgene, stained for the M-phase marker PH3 (*red*). *Scale bar, 100μm.*

(**c**) Cell cycle FACS analysis of *Actin-CreER*;*Baf53a*$^{fl/-}$ neurospheres treated with tamoxifen or vehicle and infected with empty vector or PMS factors (*Pax6*, *c-Myc* and *Sox2*). Cells were selected by a combination of blastomycin, hygromycin and puromycin prior to tamoxifen treatment. Infected cells were labeled with EdU for 2 hours for FACS analysis 72 hours after tamoxifen administration.

(**d**) Quantification of total cell numbers 72 hours after Tamoxifen treatment (normalized to condition treated with EtOH). n=2.

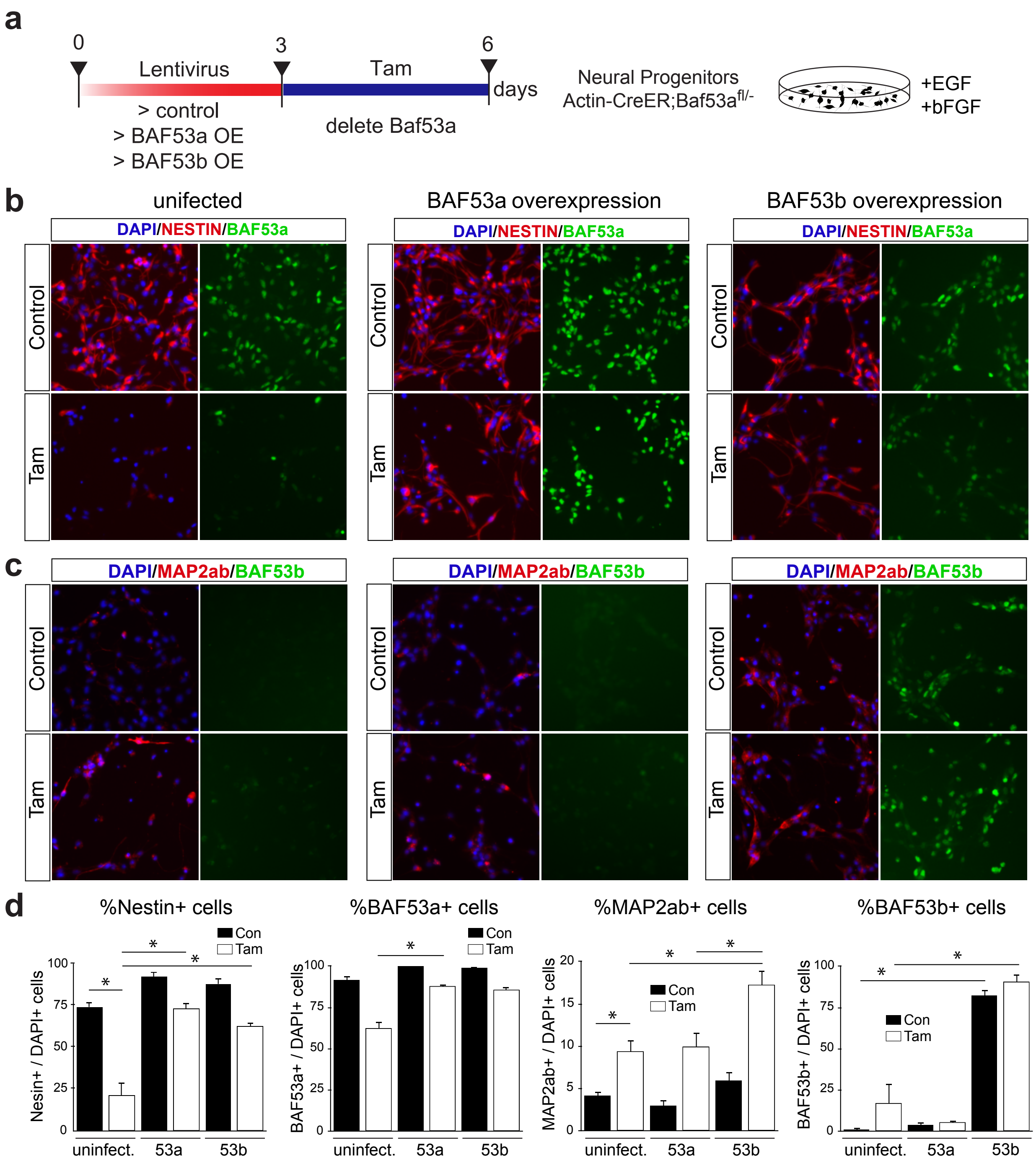


**Supp. Figure 5 (related to figure 4). BAF53b can rescue cell cycle block resulting from deletion of *Baf53a* by promoting neuronal differentiation of NPCs in vitro.**

(**a**) Experimental setup for *in vitro* lentiviral rescue experiments overexpressing BAF53a or BAF53b following deletion of *Baf53a* in NPCs.

(**b**) Immunostainings for NESTIN (*red*) and BAF53a expressing cells (*green*) in control and tamoxifen treated conditions. Nestin is a marker for NPCs and its expression is markedly reduced following *Baf53a* deletion in NPCs. Overexpression of BAF53a or BAF53b significantly increases Nestin expression following *Baf53a* deletion.

(**c**) Immunostainings for MAP2AB (*red*) and BAF53b expressing cells (*green*) in control and tamoxifen treated conditions. MAP2ab is a marker for immature neurons and its expression is detected in 10% of cells following *Baf53a* deletion in NPCs. This induced neuronal differentiation is surprising as cells are maintained in proliferative EGF/bFGF containing media. Overexpression of BAF53b significantly increases MAP2ab expression following *Baf53a* deletion.
(**d**) Quantification of cells expressing NESTIN, MAP2AB, BAF53a and BAF53b. n=3, error bars represent SEM, *$p<0.05$.

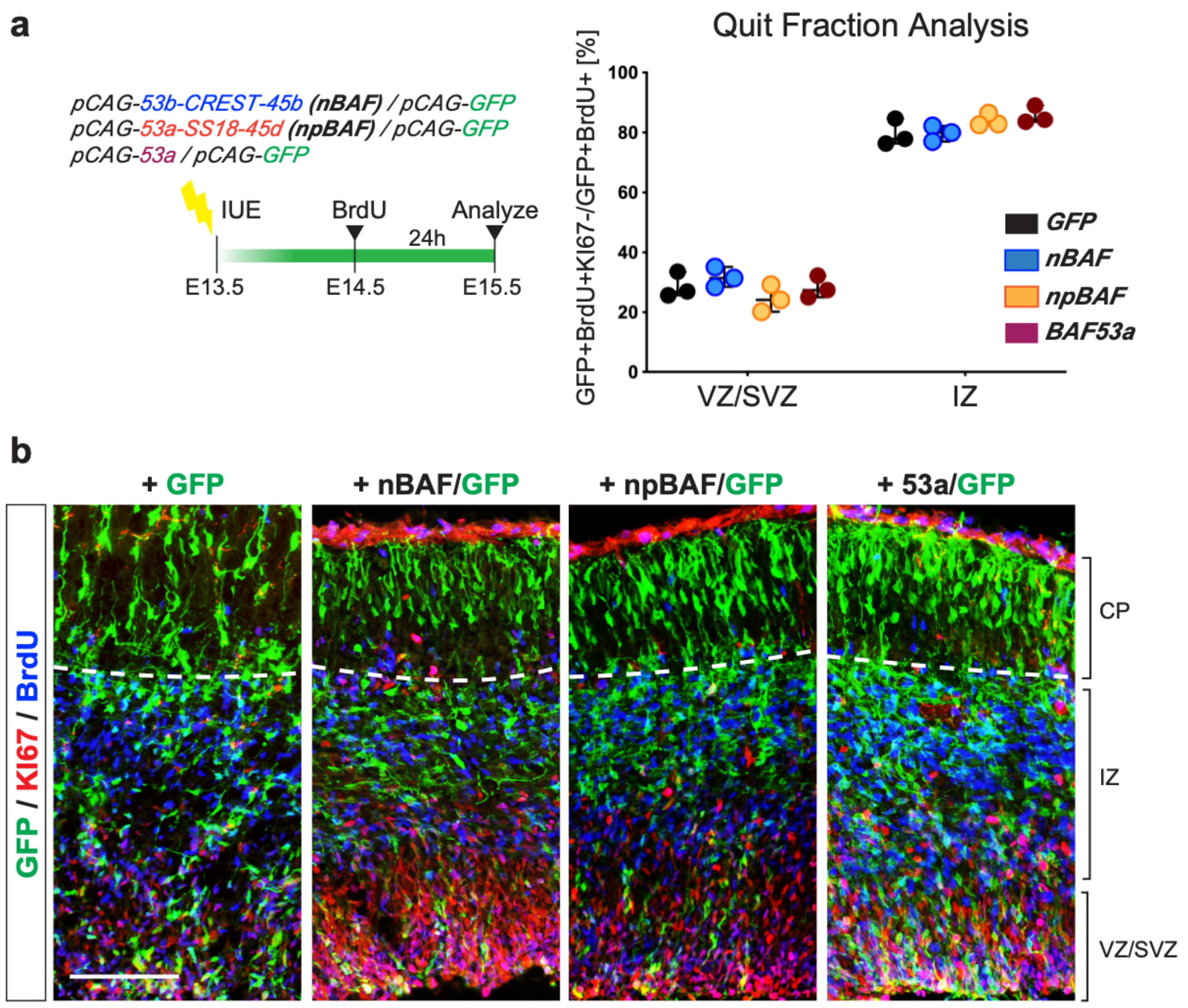


**Supp. Figure 6 (related to figure 4). Overexpression of nBAF or npBAF complex subunits, or BAF53a alone, in wild-type mice does not alter NPC cell cycle exit.**

(**a**) Schematic of experimental paradigm and expression constructs used for *in utero* electroporation of Swiss Webster timed-pregnant dams at E13.5. Quantitative analysis of the fraction of electroporated cells to have exited the cell cycle within either the ventricular-subventricular (VZ/SVZ) or the intermediate germinal zones (IZ) indicate no change in cell cycle exit. Data presented as mean +/- SEM, no significant differences observed by one-way ANOVA and post-hoc Tukey's.

(**b**) Representative images of E15.5 coronal sections from the electroporated embryonic brains used for the quit fraction analysis in (**a**) immunostained for GFP (*green*), KI67 (*red*) and BrdU (*blue*). Scale bar, 50 μm.

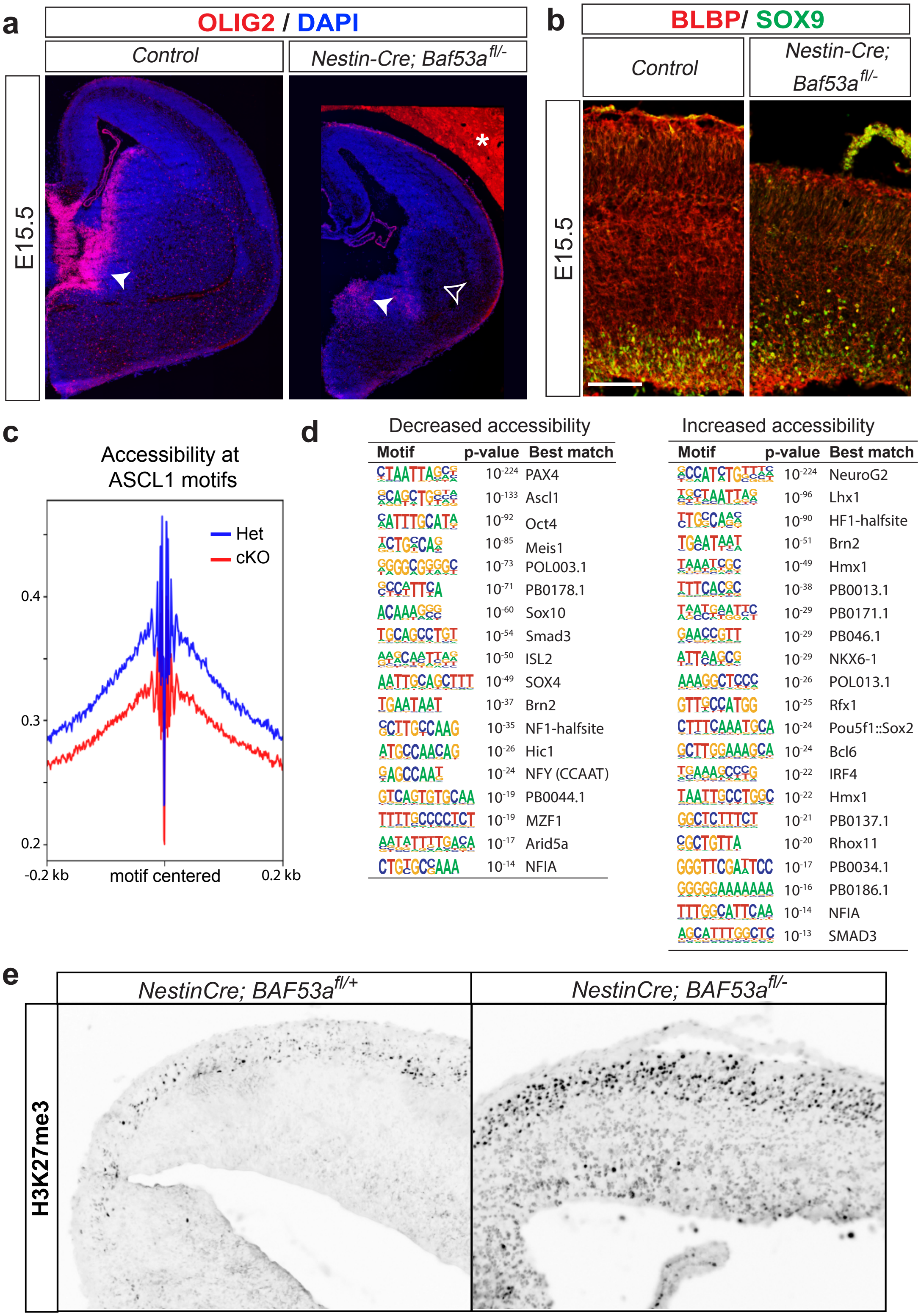


**Supp. Figure 7. *Baf53a* deletion changes the accessibility at proliferative and neurogenic gene loci.** (**a**) Immunostaining for OLIG2 (*red*) expressing cells in BAF53a mutant and control brains at E15.5. *Note asterisk (*) indicating non-specific background autofluorescent signal.*

(**b**) Immunostaining for BLBP (*red*) and SOX9 (*green*) expressing cells in BAF53a mutant and control brains at E15.5.
(**c**) Characterization of ATAC peaks at ASCL1 motifs in BAF53a mutant (cKO) and control (Het) forebrains at E15.5. Chromatin accessibility is reduced at ASCL1 motifs in mutant brains.
(**d**) Motif enrichment analysis of ATAC-seq datasets using HOMER software. Displayed are the TF motifs which are significantly enriched in peaks, which displayed either decreased or increased accessibility in BAF53a cKO brains.
(**e**) Immunostaining for H3K27me3 expressing cells in BAF53a mutant and control brains at E15.5.
